# Uncovering mechanisms of cell-free protein synthesis using tailored coarse-grained models

**DOI:** 10.64898/2026.11.24.768839

**Authors:** Christoph S. Wagner, Nadanai Laohakunakorn, Andrea Y. Weiße

**Affiliations:** School of Biology, University of Edinburgh, Edinburgh; School of Informatics, University of Edinburgh, Edinburgh

## Abstract

Cell-free protein synthesis has revolutionised molecular biology research, and it continues to be a fundamental methodology in synthetic biology today. Yet, many of its mechanistic principles remain poorly understood, limiting our ability to predict reaction behaviour. In this work, we introduce a pipeline for uncovering these principles that combines computational models with experimental validation in a design-build-test-learn cycle. First, we develop a coarse-grained ordinary differential equation model describing key processes in the reaction. Next, we adapt the model to predict system behaviour under competing mechanistic hypotheses and compare model predictions to our experimental data from batch reactions using *E. coli* extract. Our data and analysis indicate that gene length determines the rate-limiting step in transcription; it identifies transcription/translation-independent depletion of chemical energy as the primary cause for reaction termination; and finally it highlights translation as the most variable process between batches of cellular extract. Together, these results point to strategies for improving the yield and reproducibility of cell-free protein synthesis, and provide a framework for rational, model-guided design of cell-free systems.

## Introduction

Cell-Free Protein Synthesis (CFPS) is an umbrella term for methodologies that deploy the gene expression machinery *in vitro* to produce proteins, starting from a DNA template (Fig. 1A). The high throughput of and control over this methodology has made it popular for a variety of applications such as therapeutics screening, gene circuit characterisation, or molecular biosensors.^1,2^ Beyond industrial applications, CFPS also enables fundamental research into the core machinery that supports life.^3,4^ Broadly, CFPS can be divided into two categories based on the origin of their Transcription-Translation (TXTL) machinery: crude cellular extracts on one side and selectively reconstituted systems on the other. Reconstituted systems, most prominently the Protein expression Using Recombinant Elements (PURE) system,^5^ are fully determined in their composition, which allows a high degree of control and predictability. However they suffer from inefficient synthesis yields, high costs, and a complex manufacturing process.^1,2^ Extracts are powerful and relatively simple to prepare, while mimicking cellular behaviour, making them the current system of choice for industrial purposes and fundamental research into natural biology. They can be prepared from a variety of prokaryotic and eukaryotic cells, such as *Escherichia coli (E. coli)*, wheat germ, or rabbit reticulocyte.^1^ Regardless of the source organism, the crude and undetermined composition of extracts often limits reproducibility and predictability. ^2,6^ In addition to the TXTL machinery and a DNA template, CFPS reactions are powered by an energy solution that provides the enzymes with substrates (e.g. NTPs and amino acids), cofactors and a chemically favourable environment. Much work has been done on optimising this energy solution, particularly with regards to the secondary energy source, from which NTPs are regenerated.^7–10^ This study presents a workflow for dissecting the mechanisms underlying black-box extract systems, to enhance their predictability.

In a batch reaction, CFPS occurs following a universal three-phase pattern (Fig. 1B): an initial lag-phase, caused by the time required for transcription, translation, and folding/maturation of a protein of interest, is followed by a linear production phase, the effective rate of which is determined by the rates of the individual comprised steps, until the reaction finally terminates. *In silico* models can be used to predict the behaviour of a system under untested conditions^11^ (Figs. 1A and B). In particular, mechanistic models, which attempt to describe a behaviour by making assumptions about the underlying processes, can yield understanding of the system’s operations. ^11^ Naturally, the detail of understanding is limited by the granularity, i.e. level of detail, of the model. In this work, we build a coarse-grained model, leveraging its simplicity and interpretability. We then modularly add mechanisms to contrast different hypotheses and their agreement with data (Fig. 1B). In this way, the model can help us gain a mechanistic understanding of the system under study.

**Figure 1:**
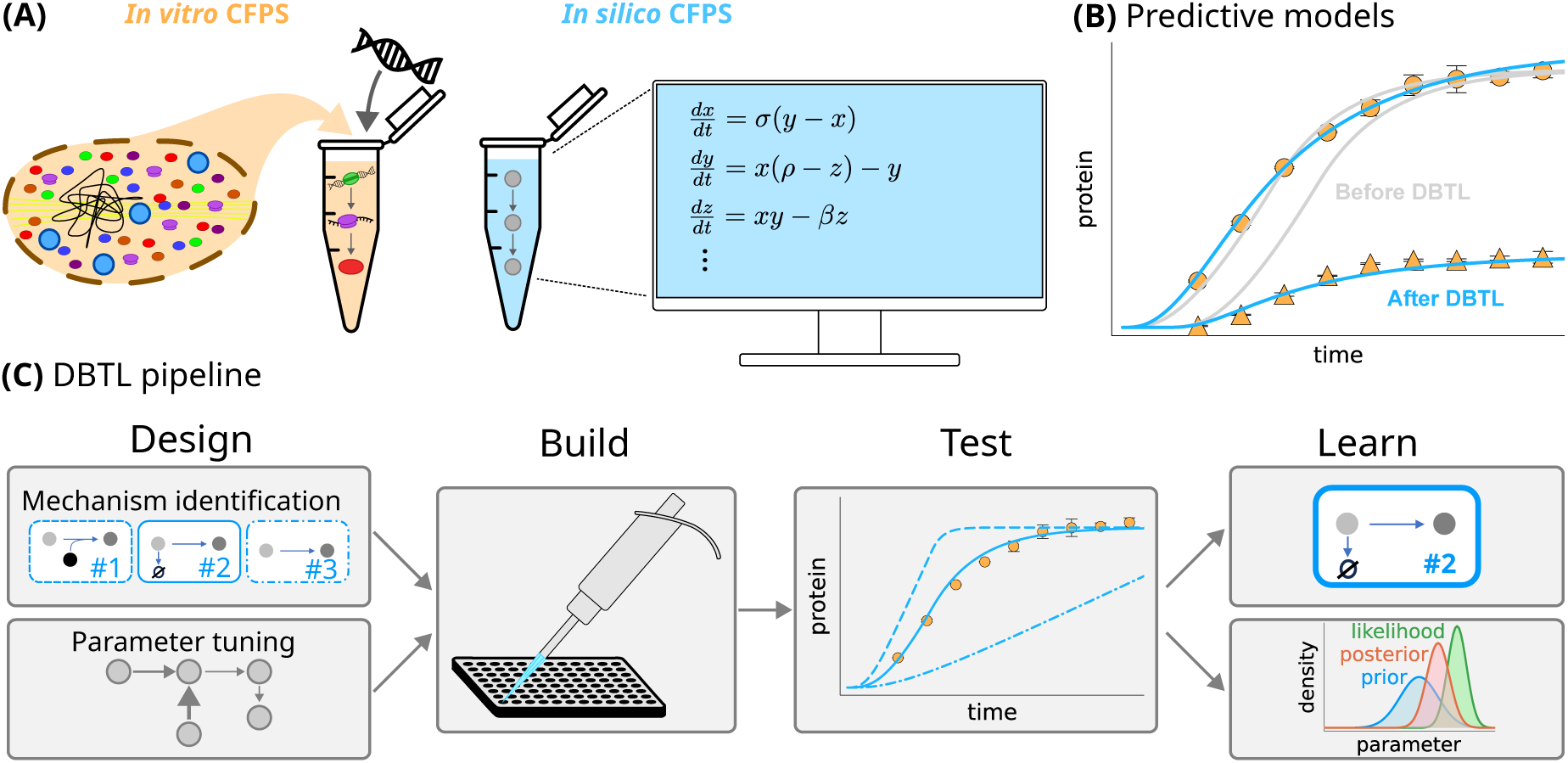
Developing a model of CFPS by iterative comparison with and adaptation to experimental data. **(A)** Ordinary differential equations can mirror *in vitro* CFPS reactions, predicting properties like rate, lifetime and yield of the *in vitro* reaction. **(B)** Il-lustration of model improvements during a Design-Build-Test-Learn (DBTL) iteration: An initial model (grey lines) captures one dataset (circles) but not another (triangles). After DBTL, the adjusted model (blue lines) captures both datasets. **(C)** The presented DBTL pipeline facilitates model adjustments by designing and building an experiment, comparing model predictions to experimental data and identifying the required changes to align the two. Mechanism identification or tuning of parameter values (top and bottom, respectively, in ‘Design’ and ‘Learn’) are two possible applications of the pipeline.

Such iterative rounds of improvement, or tailoring, are central to the DBTL principle (Fig. 1C): a question is posed (‘Design’) and the corresponding model and experimental setup developed (‘Build’). Experimental data is gathered (‘Test’), which is finally used to answer the initial question (‘Learn’). This principle is highly flexible and has been used in the context of CFPS previously.^12,13^ It may be applied to a variety of questions, such as qualitative selection of model mechanisms between competing hypotheses or quantification of a particular model parameter. Furthermore, DBTL may be deployed in iterative cycles to deepen the understanding of one pathway or leveraged non-consecutively as a pipeline to explore various pathways of one system. ^13^

In this work, we present a baseline coarse-grained model of *E. coli* extract CFPS, which recovers the typical lag-production-termination behaviour described above. We then explore cases the model fails to accurately describe, uncovering mechanistic limitations of model assumptions. We first seek to characterise the rate-limiting steps in transcription and translation, thereby revealing how their kinetics may be engineered. We then determine the mechanism behind reaction termination and explore resource depletion over time, making reaction longevity more predictably and tunable. Finally, we investigate the source of batch-to-batch variability, a major roadblock in reproducibility of lysate-based technology. To answer each of these questions, we start with a baseline model that we selectively extend until experimental data is accurately recovered, which yields insights into mechanisms required to capture the complexity of the data. Beyond improving model predictability, these insights point to strategies for improving the yield and reproducibility of CFPS, and provide a framework for model-guided design of cell-free systems.

## Results

### A coarse-grained model of CFPS that recovers lag, production and termination phases

We built a baseline model of CFPS describing core underlying processes (Fig. 2). Based on a system of ordinary differential equations the model describes the central dogma of molecular biology: DNA *d* is transcribed into mRNA *m*, which binds the ribosome *r* to form the translating complex *c* and translate protein *p*, using up energy *e* in the process. The protein finally matures into *p_m_*. We describe transcription of mRNA *m* and translation of protein *p* as single-step reactions with maximal effective rates determined by rate constants *k_tx_* and *k_tl_*, similar to previous works. ^14–16^ Their rates are assumed inversely proportional to the product length *n_m_* and *n_p_*, respectively, to account for elongation, and depend on the availability of a generic “energy” species *e* in a saturable manner. ^17^ Finally, both transcription and translation are catalysed by their respective enzymes, i.e. RNA Polymerase (RNAP) and ribosomes, respectively. Transcription is performed by RNAP on a DNA template, both of which exit the reaction unaltered. Effective rates are derived as follows.

We assume that the concentration of T7 RNAP is abundant compared to the supplied DNA concentration (on the order of 10 µMT7 RNAP and up to 5 nM DNA, see SI), making DNA *d* the effective enzyme and as such the transcription rate *v_tx_* can be written as (see Weiße et al. for a derivation)

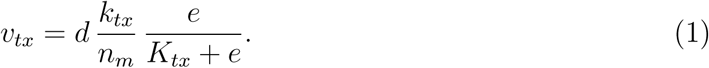

Our estimations placed the concentration of ribosomes *r* on roughly the same order of magnitude as *m*, since we added ribosomes at around 500 nM, while mRNA is typically produced on the order of 10-100 nM.^14,18^ Hence, neither could be assumed abundant. To account for competitive binding, we modelled *r* and *m* to undergo a diffusion-limited, reversible binding step to form the translation complex *c* with rate constant *k_b_*, which can dissociate with rate constant *k_u_*, respectively, according to the law of mass-action.^17,19^

To avoid complex modelling of polysomes, we introduced a scaling term *p_b_* to account for formation of several complexes *c* per ribosome-mRNA binding event by

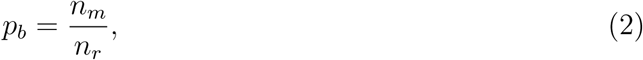

where *n_r_* = 270 is the average length of mRNA occupied by one ribosome.^20^ Hence, the simplifying assumption is made that each mRNA binds *p_b_* ribosomes upon a binding event, removing one mRNA and *p_b_* ribosomes from their free pools to form *p_b_* complexes *c*. After this, *c* is the species that catalyses translation, releasing protein alongside free mRNA and ribosome with rate

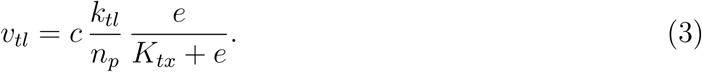

In our experimental setup we use the fluorescent reporter protein mCherry, which matures after translation, and only then becomes observable.^21^ We modelled this as a first-order reaction with rate constant *k_m_*, giving *v_m_* = *k_m_ p* (see SI for more details), and producing

**Figure 2:**
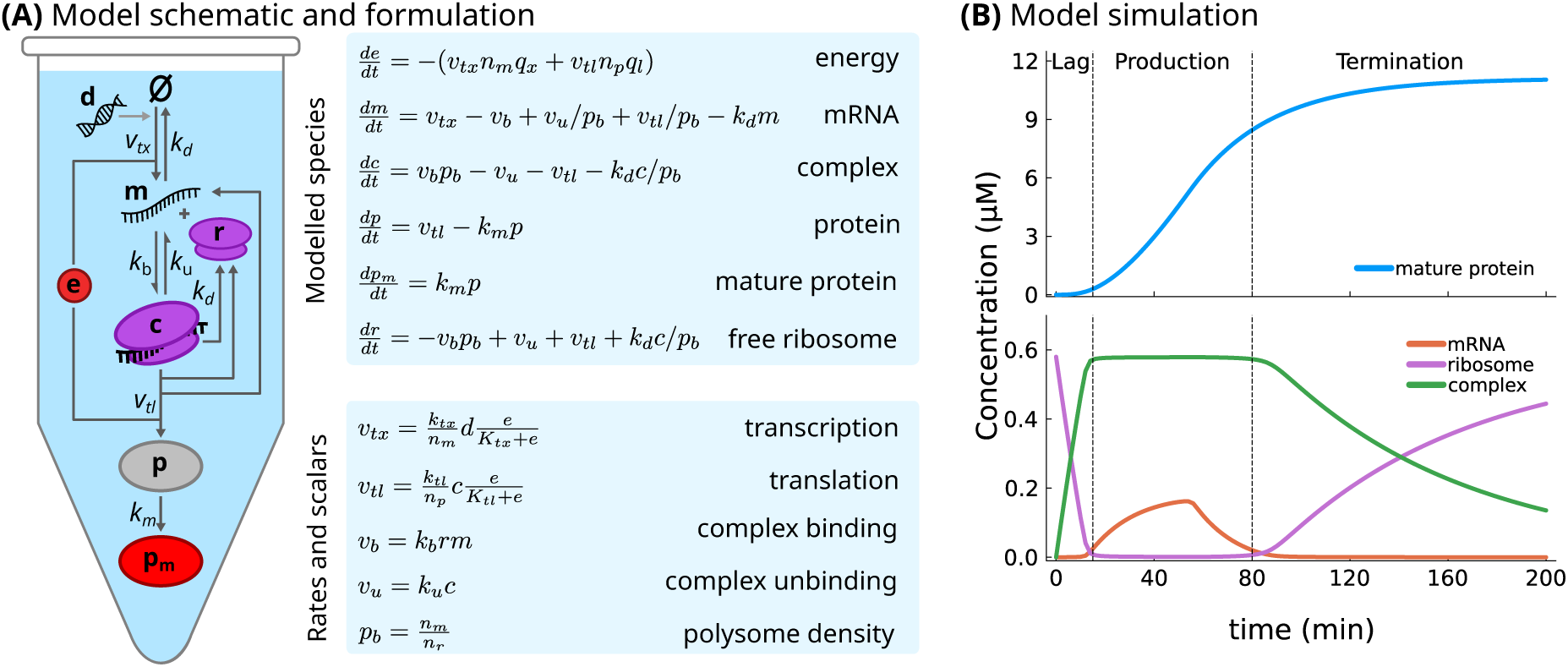
Baseline CFPS model used in this study. **(A)** A schematic of the modelled pathways is shown on the left with species in bold, and rates in italic. The rates of change for each species and rate terms are listed on the right: mRNA *m* is transcribed from DNA *d* at rate *v_tx_*. It then dynamically binds ribosome *r* to form the translating complex *c*, which translates *p* at rate *v_tl_*. *p* finally matures into *p_m_*. Transcription and translation consume energy *e*. **(B)** A simulation of the model with default parameterisation (Tab. S1) is shown on the right, split between the observable fluorescent protein (top) and background species mRNA, ribosome, and translation complex (bottom).

*E. coli* extract contains RNases, which degrade mRNA.^22^ The dominant degradation pathway is internal cleavage by endonucleases, with degradation from the ends through exonucleases playing a secondary role.^23^ As the ribosome density on mRNAs in cellular extract is relatively sparse at around 270 nt between ribosomes^20^ and ribosomes only protect their footprint from degradation through endonucleases,^24^ we assumed free and bound mRNA susceptible to degradation by the same rate constant *k_d_* in a first-order kinetic reaction.

For simplicity, we modelled both transcription and translation to consume the same enegetic substrate *e*. While the two processes depend on various and partially non-overlapping resources, both require NTPs to take place. As such, “energy” can be thought of as the chemical energy stored in each phosphodiester bond in NTP. Each elongation step of the RNAP and ribosome consumes around *q_x_* = 2 and *q_l_* = 4 phosphate bond equivalents, respectively.^25^ We therefore set the consumption of energy through transcription and translation negatively proportional to their respective rates, the length of their products, and the phosphate bond equivalents per elongation step.

The full model formulation alongside estimated default rate parameters, carefully collated from the literature, and initial species concentrations can be found in Tab. S1. It recovers the three phases of CFPS (Fig. 1B), with the lag phase dominated by the maturation rate *k_m_*, the production phase determined by the combination of transcription, translation and maturation, and termination caused by the depletion of *e*.

### Gene length determines rate-limiting step for transcription while elongation primarily limits translation

In the baseline model described above, mRNA and proteins are synthesised at rates inversely proportional to their product lengths to account for the sum of elongation steps necessary to produce the full-length polymers. However, polymerisation is generally a process comprised of three phases: initiation, elongation and termination. During initiation, the enzyme binds the template at the promoter or ribosomal binding site and the complex may undergo a series of intermediate states before forming a stable elongation complex. ^26–33^ During elongation, the complex repeats the same step *n_x_* times. Finally, polymerisation terminates at the terminator or stop codon and the enzyme releases the template.

There are different ways of modelling such multi-step processes. Each step may be defined explicitly, which enables the model to capture intricate effects but requires comprehensive data for parameterisation.^19,34^ To balance complexity with predictive power, a multi-step process may alternatively be described by the slowest, and thereby rate-limiting step, disregarding the other steps as negligibly fast.^14,16,35,36^ This reduces the number of parameters in a model^19,37^ and facilitates transfer of fundamental information between studies.^38^

To identify the rate-limiting steps in transcription and translation in the modelled system, the respective individual rates may be compared. Transcription has previously been suggested to be initiation-limited in T7 RNAP-controlled extract-based CFPS.^34^ The slowest experimentally estimated T7 RNAP initiation rates on the T7 *ϕ*_10_ promoter are around 0.3 min*^−^*^1^,^32,39–41^ while the fastest published elongation rates are around 200 nt min*^−^*^1^.^41–43^ For transcripts of few hundreds of nucleotides length, as is typical for CFPS, neither initiation nor elongation may therefore safely be assumed the rate-limiting step. In contrast, most experimental evidence points towards elongation as the rate-limiting step in translation in *E. coli* CFPS,^20,34,43–46^ however, this is also occasionally challenged.^47^ To the best of our knowledge, the contributions of the individual steps to the overall transcription and translation rates in *E. coli* CFPS have never been measured explicitly and are therefore still an open question.

To approach this issue, we made two simplifying assumptions. First, we assumed that initiation and termination rates depend exclusively on their respective template sequences and the enzyme in question. This means that constructs with the same initiation and termination sequences should have constant initiation and termination rates, which can be further condensed into one lumped initiation and termination rate. Since initiation is usually considered slower than termination,^41^ we will be referring to this lumped constant rate as the initiation rate in the following (Fig. 3B). The second assumption is that the effective elongation rate scales primarily with the length of the product and that any secondary effects, such as those of sequence composition or template secondary structures are negligible against the sequence length^48^ as well as constant for repeated sequences.

If a process is purely initiation-limited, it should take place at the same rate for varying lengths, while an elongation-limited process should vary with gene length. Accordingly, we prepared six DNA constructs that would allow us to decouple the initiation from the elongation rate, by varying the gene length, while keeping the initiation and termination sequences constant (Fig. 3A). We modified a plasmid carrying mCherry-Broccoli under T7-control^35^ by either inserting copies of a mutated, non-fluorescent ‘dead’ mCherry, with a basepair change leading to an M71E mutation,^49,50^ exactly after the 3’ end of the original start codon, or by removing a portion or all of the coding sequence. This produced six transcription templates with constant promoter and terminator sequences but sequences of varying length ending in a Broccoli RNA reporter,^51,52^ which could be used for deconvoluting the effective transcription rate (Fig. 3A, green). Within this set, three constructs further contained identical ribosomal binding sites followed by coding sequences of varying length, ending in mCherry reporter, which could be used for deconvoluting the effective translation rate (Fig. 3A, red).

We then carried out CFPS reactions using each of the constructs, and collected Broccoli as well as mCherry signal as time series data (Fig. 3C). We measured data for the three mCherry constructs twice on separate occasions. After identifying the linear production phases in a window, set visually between 6-25 min for transcription and 80-150 min for translation, we performed a linear fit to determine the slope, i.e. the effective rate (Fig. S19).

Next, we modified the model to determine which hypothesis best explains the data (Fig 3D). We substituted the rate constant term *k_tx_/n_m_* in Eq. (1) with a constant initiation rate *k_txI_* (initiation-limited regime), kept it the same (elongation-limited regime), or extended it to include contributions from both initiation and elongation by means of partition analysis.^53^ Under all three regimes, we then inferred *k_txI_*, *k_txE_* or both by Bayesian inference (see SI), by simulating the model, extracting the slope of the simulation and minimising the distance between experiments and simulation of this surrogate term. ^6^ For full details on Bayesian inference, refer to the methods and SI (Figs. S5-S13).

**Figure 3:**
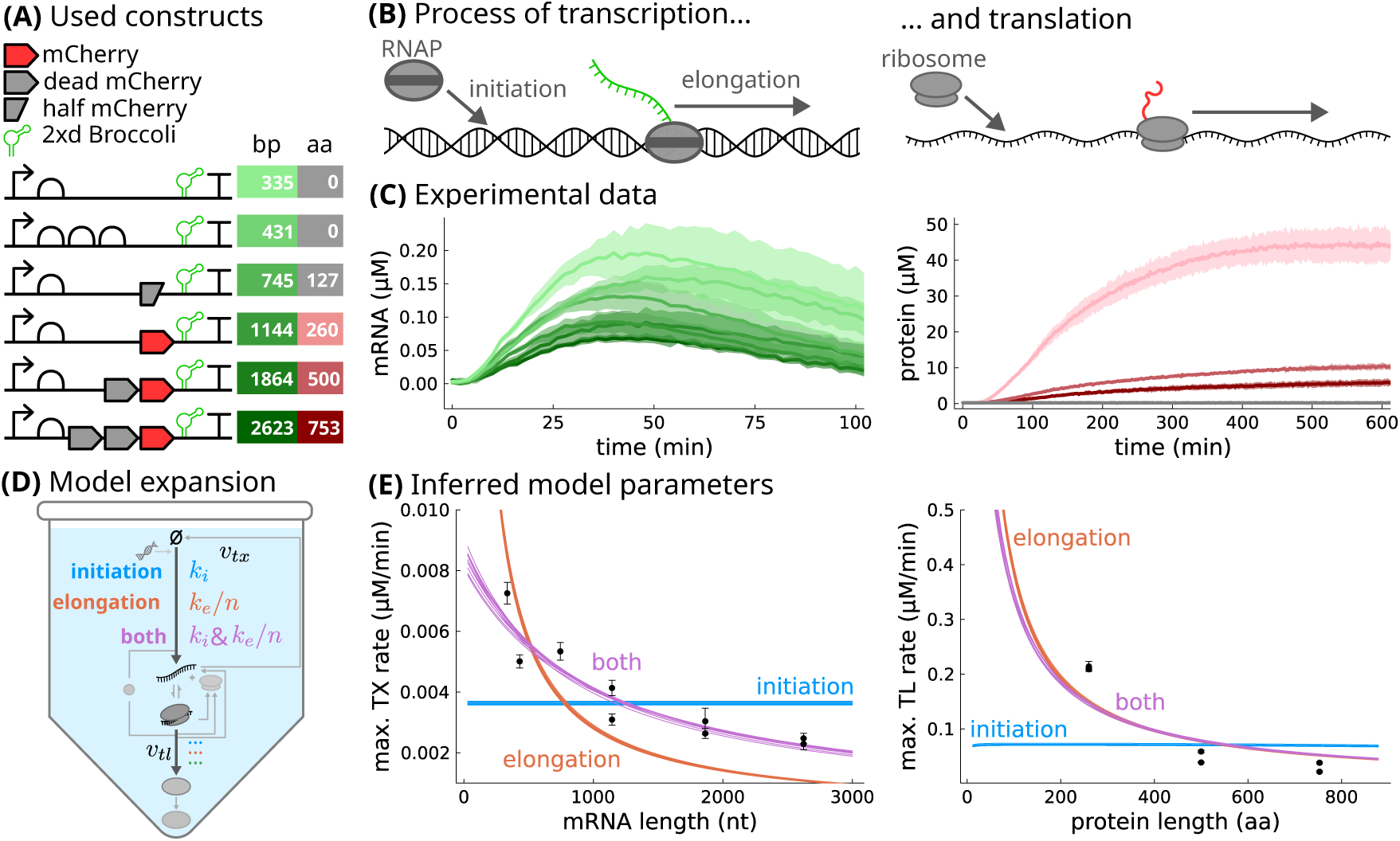
Rate-limitations in transcription and translation. **(A)** DNA constructs cloned for this study. The graphic follows Synthetic Biology Open Language conventions, with unique symbols annotated above. Transcript and protein lengths are highlighted in green and red, respectively. Numbers in grey are not observable experimentally. **(B)** Studied processes: initiation and elongation in transcription (left) and translation (right). **(C)** Fluorescence signals for transcription (Broccoli, left) and translation (mCherry, right) observed for the constructs shown in (A), with shaded areas showing one standard deviation of the average over triplicates (solid line). Colours correspond to the construct lengths in (A). **(D)** Investigated model extensions, implementing initiation-limited (*k_i_*, blue), elongation-limited (*k_e_/n*, orange) or both limitiations (purple) for transcription or translation. **(E)** Maximal rates of transcription (left) and translation (right) against construct lengths. We compare observed rates (black circles as mean of three samples with one standard deviation as errorbar) from data in (C) to fitted model predictions (lines) under the hypotheses depicted in (D). For each hypothesis, we sampled 10 parameter values from the posterior to simulate the model and plot the respective maximal rate against the underlying product length (10 lines coloured according to D).

We found that the transcription data can be best explained when taking contributions from both initiation and elongation steps into account (Fig. 3E, left). Assuming an exclusively initiation-limited regime, maximal transcription rate should remain constant with varying gene length. However the data shows a decrease in rate with increasing length. Under the assumption of exclusively elongation-limited transcription, the model predicts a stronger effect of length on rate than is observed. However, if both contributions are taken into account, the model is able to accurately recover the experimental data. By plotting the individual contributions of initiation and elongation, respectively, on the total transcription rate under the combined regime, we find that the inferred initiation rate of 16.13*±*0.65 min*^−^*^1^ dominates for genes shorter than around 1000 nt, while longer genes are dominated by the inferred elongation rate of 14356 *±* 662 nt min*^−^*^1^ (Fig. S20a). We therefore concluded that in this system and for the studied gene lengths, both initiation and elongation are important to capture the overall transcription rate.

We carried out the same analysis for translation, now modelling transcription under the combined regime (Fig. 3E, right). Again, an exclusively initiation-limited regime was not able to recover the experimental data. The elongation-only regime aligned nearly perfectly with the combined regime and after inference the combined regime was dominated by the elongation rate as well. This can also be observed when plotting the individual contributions of the combined translation rate separately (Fig. S20b). This result aligns with the assumption that translation in CFPS is elongation-limited.^20^

However, it became apparent that the data showed an even stronger dependence of translation rate on gene length than could be recovered in the elongation-only regime. The translation rate of the shorter gene was underestimated and that of the two longer genes overestimated by the model when compared to the experimental data (Fig. 3E, right). Previous work has suggested that protein truncation is a major effect in CFPS, both in extract^54,55^ and in reconstituted systems.^15,56^ Alternatively to the simulations in Fig. 3E, a model adaptation that takes truncation into account on top of the competing rate contributions was able to more accurately recover the experimental data, suggesting one potential explanation for the observed divergence (SI, Figs. S11-S13, S20).

We finally concluded that, within the range of gene lengths tested, transcription rate is limited by both initiation and elongation, while translation rate is to a large extent limited by elongation rate, noting that other untested phenomena, such as protein truncation, may also contribute.

### CFPS reaction lifetime is primarily limited by a mechanism independent of TXTL activity

We next examined limitations to reaction lifetime in batch CFPS. There are various possible reasons for reaction termination, which may be classed as (1) TXTL-dependent or (2) TXTL-independent (Fig. 4A). In the first case, a reaction is stable over time, proceeding for as long as the required resources are present and functional, and terminating once a limiting resource is depleted or otherwise unavailable. In the second case, reactions are unstable over time, with lifetimes limited by a factor unrelated to their own activity.

Both TXTL-dependent^57,58^ as well as independent^5,7,59–61^ termination of CFPS have been described in the literature, which indicates that the mechanism may depend on the exact CFPS system in question. Indeed, the results for termination of PURE differs from those indicated above.^15,59^ Our default model implements termination upon complete consumption of *e*, i.e. a TXTL-dependent mechanism, however this may not be accurate. To be able to predict system behaviour under various conditions, we seek to identify the contributions of TXTL-dependent and -independent processes on reaction termination.

To identify the correct termination category, we first simulated varying amounts of substrate available to the reaction (i.e. the initial energy *e*_0_), as well as the rate of a reaction, through varying the ribosome concentration *r*_0_ and DNA concentration *d*. We implemented TXTL-independent termination in the model by introducing an energy degradation rate *k_de_*. However, other decay pathways such as those of ribosomes and DNA, produce qualitatively similar results. Under this assumption, both varying substrate availability or ribosome concentration impacts the final yield as well as the maximal rate of the reaction (Fig. 4B, top). On the contrary, the default model without such a degradation rate assumes that a reaction is terminated exclusively upon TXTL-dependent consumption of resources. In this case, varying substrate availability only affects the final yield but not the rate, while varying enzyme activity only affects the rate while the yield remains constant (Fig. 4B, bottom).

Two characteristic properties of a batch CFPS reaction are its maximal rate and final yield (Fig. 4C). They can be represented as a point on a 2D-plane, where the yield is plotted against the rate. A series of experiments can be represented as a trajectory on that plane. By plotting yield and rate normalised to their respective maxima per series, we can examine the trade-offs generated by the different scenarios described above (Fig. 4D). Under the hypothesis of TXTL-dependent termination, increasing enzyme activity increases the rate while yield remains constant, thus resulting in a horizontal line along the top edge. *Vice versa*, varying the substrate availability increases yield at constant rate, resulting in a vertical line along the right edge. On the contrary, if TXTL-independent resource degradation is the cause of termination, then an increase in either substrate availability or enzyme activity leads to a diagonal line, where yield is directly correlated with rate, as more substrate can be converted into product before it is degraded. A line orthogonal to the diagonal is theoretically also possible but has rarely been described and will therefore not be further discussed here (Fig. S21).

**Figure 4:**
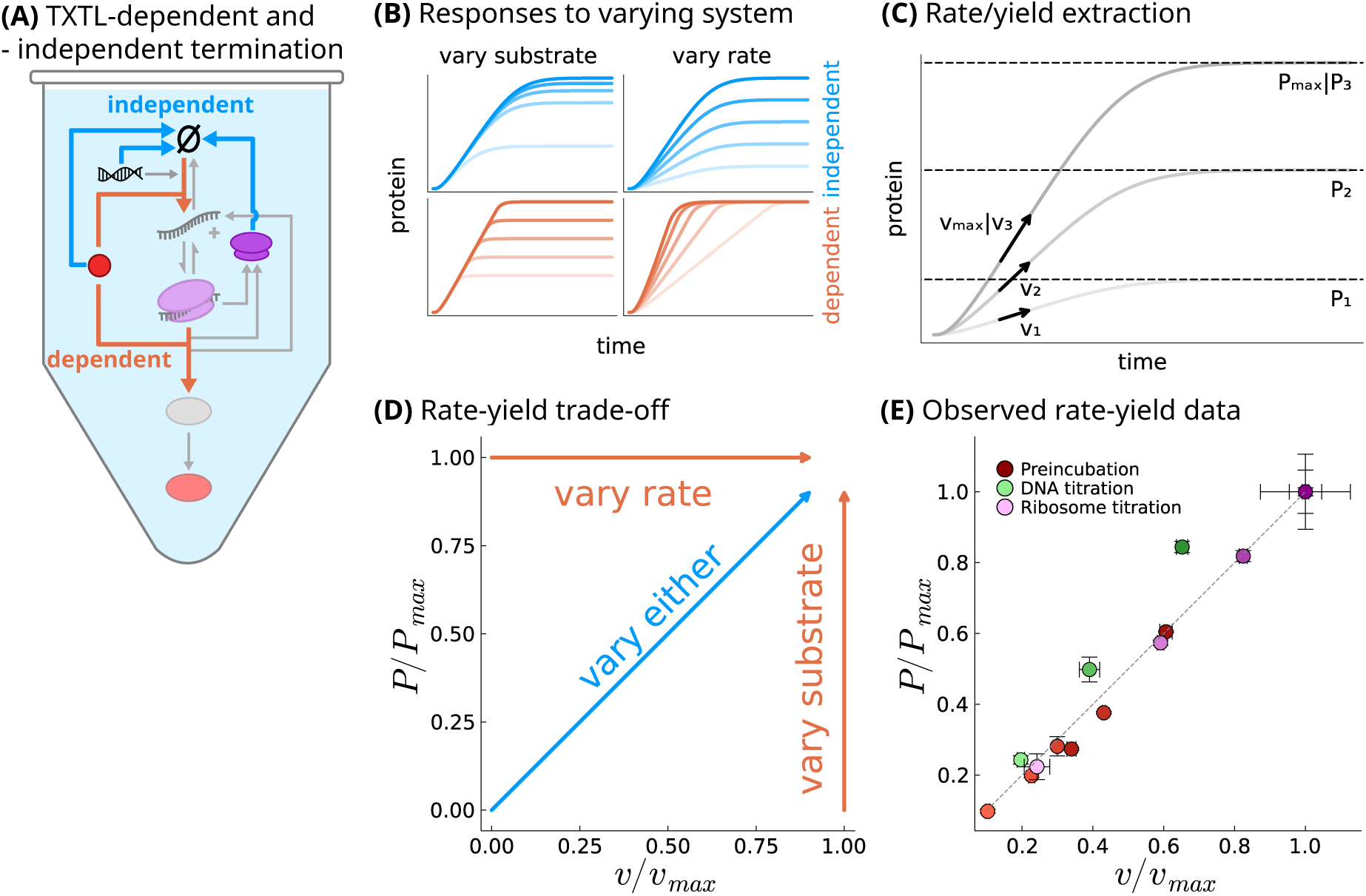
Termination of CFPS. **(A)** Two possible categories of reaction termination may be implemented in the model: TXTL-independent (blue) or TXTL-dependent termination (orange). **(B)** Depending on the termination mechanism in (A), a reaction shows different behaviour upon manipulation of its parameters. **(C)** The rates and yields of a series of reactions are extracted as their maximal slopes and protein concentrations. They can be normalised to 1 by dividing by *v_max_* and *P_max_*, respectively. **(D)** The rates and yields of each experiment, normalised to the maximum value for each dimension and plotted against each other, reveals qualitatively different trajectories between the hypothetical termination mechanisms and manipulations from (B). **(E)** Data from three different experimental assays, processed according to (C) and plotted as in (D), lie on the diagonal (*y* = *x*, dotted line). Dark shading indicates shorter preincubation without DNA, more DNA and more ribosomes, respectively, errorbars denote one standard deviation in either dimension. The ‘Preincubation’ assay was measured with duplicates, the other two with triplicates.

We then conducted multiple series of experiments, at different maximal reaction rates, by varying the concentration of DNA or ribosomes in the CFPS reaction. Separately, we varied CFPS preincubation time before addition of DNA between 0-60 min, to check for stability over time (Fig. S22). We extracted normalised rates and yields of each series according to Fig. 4C and plotted them in rate-yield space (Fig. 4E).

Our data strongly suggest that the analysed system is terminated partially independently of TXTL, as most points lie on the diagonal. However, titration of DNA shows a slight deviation, as the highest tested DNA concentration produces a slightly lower yield than would be expected from its rate, indicating that a saturation point may have been reached, that had not been accounted for in the model. With the prospect of TXTL-independent resource degradation, we next asked what resource is affected.

### Degradation of chemical energy terminates reactions

Multiple causes for TXTL-independent resource decay have been discussed and identified previously, which can be categorised into four possible scenarios (Fig. 5A).

First, reactions may run out of chemical energy and substrates.^5,7,19,59–62^ Chemical energy can be thought of as NTPs and their precursors. Since *E. coli* extract contains an uncharacterised pool of enzymes, it is likely that the TXTL machinery is not the only consumer of NTPs. Alternatively, TXTL substrates, including NTPs and amino acids, may also be funnelled through non-productive pathways, depleting the reaction of its building blocks.^7,58–61^

**Figure 5:**
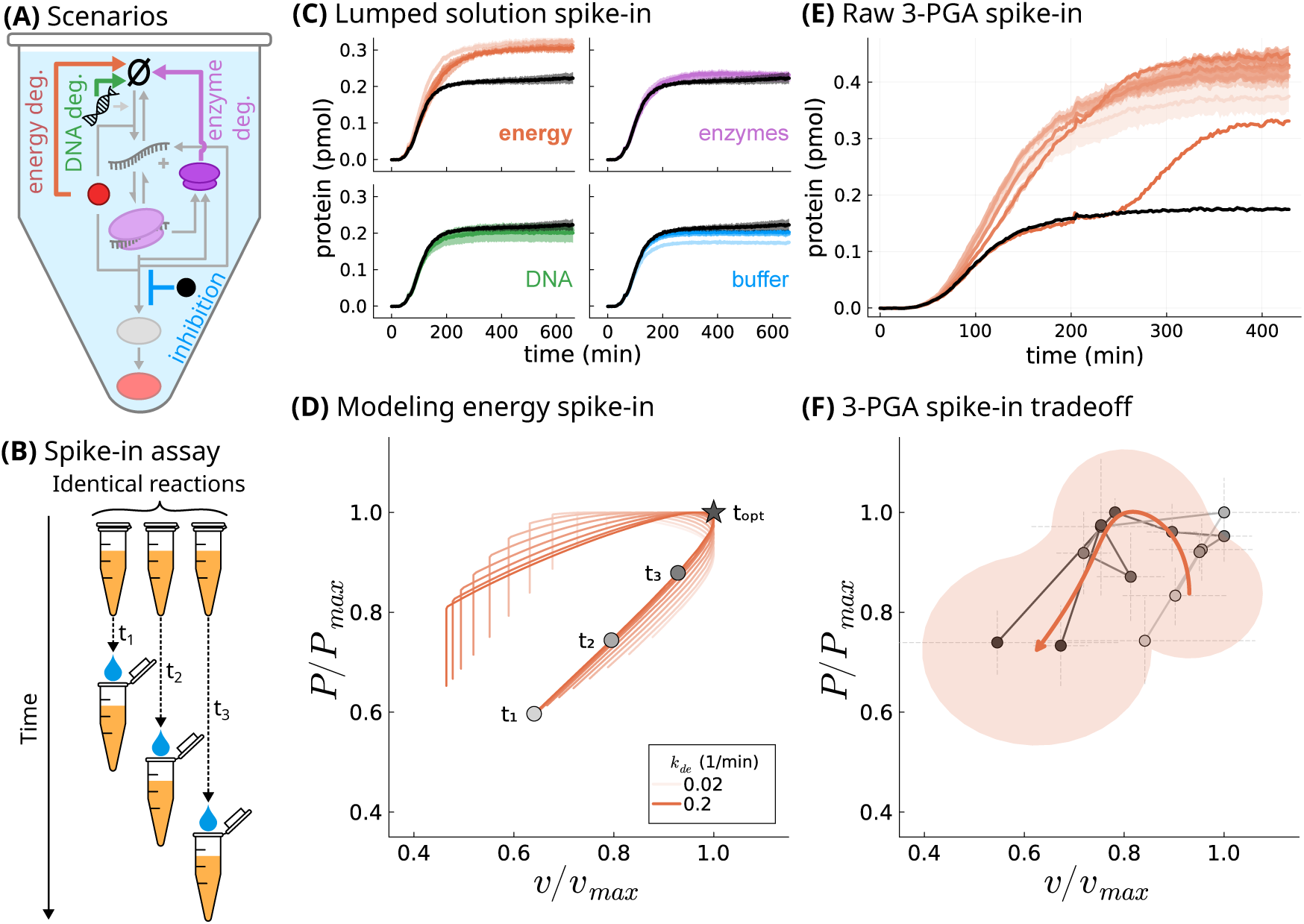
Characterisation of resource degradation dynamics. **(A)** We considered four hypothetical scenarios of TXTL-independent termination: energy/substrate degradation (orange), enzyme degradation (purple), DNA degradation (green), inhibitor accumulation (blue). **(B)** Spike-in assay developed to probe resource degradation over time. Identical reactions are started simultaneously and spiked individually over time with a reagent of interest. **(C)** Results of spike-in assay testing the broad reagent categories from (A): energy solution, lysate, DNA and buffer after 20, 40 or 60 min (colour intensity). Fluorescence of mCherry over time is displayed as the average value (solid line) with one standard deviation (shaded region) over triplicates. **(D)** Model prediction for spike-in of *e* over a time period of 200 min (‘Addition time’) for *k_de_* ranging between 0.02-0.2 min*^−^*^1^ (colour intensity), plotted in rate-yield space. An ideal spike-in time maximises rate and yield (star) **(E)** Raw results for 3-Phosphoglycerate (3-PGA) spike-in between 0-50 min (lighter orange) or after 180 min (dark orange), compared to no spike-in (black). Fluorescence data as in (C) with duplicates where a shaded area is indicated. **(F)** Rate and yield properties from two experiments like the one shown in (E) plotted in rate-yield space, where light grey indicates early addition and dark grey late. The orange arrow indicates a gaussian process regression fitted to the data with shaded regions indicating one standard deviation in both dimensions as ellipses around each curve point. Experimental uncertainty for single measurements was approximated by the largest uncertainty in the respective assay.

Second, TXTL enzymes may be slowly degrading, which would eventually halt a reaction, even if energy and substrates are still available. This has been suggested previously^58^ but seems more pronounced in PURE than in extract.^15^

Third, DNA may be unstable in extract. Since this study uses plasmid DNA, this scenario is unlikely, as plasmid DNA has, to our knowledge, not been observed to degrade in such an environment to an observable degree. However, we considered the scenario hypothetically possible for this work.

Fourth, the reaction environment may experience metabolic changes that slow down and stop TXTL. Various effects have been described previously, including depletion of magnesium,^15,63^ accumulation of inhibitors such as inorganic phosphate ^7,19,41,57,59,60^ or macro-molecular crowding.^6^ For the purpose of this work, we considered all of these under the naϊve mechanism of ‘accumulating inhibitors’.

We reasoned that one or more of the above scenarios may have an effect and that the magnitude of their effects may vary over reaction time. To elucidate the contributions of each scenario and their time-dependence, we devised a fed-batch experiment (Fig. 5B): We prepared a set of identical reactions simultaneously. Then, we periodically spiked a different subset of reactions with fresh energy solution, lysate, DNA or buffer, each corresponding in that order to the four above scenarios. The hypothesis was that the reagent leading to an increase in rate or yield in a reaction is likely to be the limiting reagent at that time.

Spike-in of enzymes, DNA and buffer had negligible effects on reaction behaviour, whereas energy solution increased final yield similarly at all tested spike-in times, and rate only when added early (Fig. 5C). This aligned well with previous research suggesting fuel or substrate degradation.^5,7,58–61^ However, the energy solution contains 16 components that cover all possible aspects of the reaction. We therefore perfomed a second, single time-point spike-in experiment to test the most likely candidates within the energy solution (NTPs, 3-PGA, amino acids and magnesium), which identified NTPs and 3-PGA as the first limiting resources (Fig. S23). Contrary to previously studied systems,^7,58–61^ our results indicate that the reactions described here are limited by chemical energy and its precursor, which power both transcription and translation directly.

So far, we showed that TXTL-independent resource decay, including decay of chemical energy players, affects the duration of a reaction. This indicates that higher initial concentrations of NTPs or 3-PGA do not necessarily lead to the highest possible conversion efficiency of substrate to product. We therefore next asked if the time at which additional energy is spiked-in may thus be an amenable tuning knob to improve efficiency.

Introducing a first-order energy decay term, dependent on energy degradation rate constant *k_de_*, we simulated energy spike-in at various times and for varying decay rates. Fig. 5D shows how rate (horizontal axis) and yield (vertical axis) of a reaction benefit from spike-in of energy, depending on spike-in time (position along the line starting at t_1_) under different assumptions for the energy degradation rate constants. The model predicted that there exists an optimal addition time, that leads to the highest gain in both rate and yield, at the turning point of the curve (star in Fig. 5D). This turning point coincides with the time of depletion of the original energy pool *e*_0_, which we arbitrarily set as *e <* 2000 µM (Fig. S24).

If energy is added to a running reaction before *e*_0_ is depleted, excessive energy is wasted by decay (Fig. 5D, upwards branch). Otherwise, if energy is added too late after depletion of *e*_0_, other TXTL resources such as *m* or *c* are wasted by degradation and must be synthesised again after energy spike-in (Fig. 5D, downwards branch).

Encouraged by the model predictions, we conducted a series of experiments spiking 3-PGA spiked into a running reaction at different time points and monitoring resulting benefits to rate and yield (for an example experiment, see Fig. 5E). We extracted normalised rates and yields for each sample and plotted them in rate-yield-space to compare with model predictions (compare Figs. 5D and F). Indeed, we observed an optimal addition time of 3-PGA at around 30 min after onset of the reaction. Addition before this time displayed lower yields and slightly lower rates (upwards branch), while later addition mostly reduced rate (downwards branch), until addition became so late that production was biphasic (Fig. 5E) and no rate benefit could be observed over the control sample (vertical line portions in Fig. 5D).

These results indicate that chemical energy is a dominant limiting factor in the system studied here and that its decay dynamics imply an optimal timing for supplementing a depleted reaction with fresh energy for extracting the highest yield. Using models to fine-tune batch reaction conditions over time is gaining traction as a useful technique for optimising such reactions e.g. for industrial purposes.^12^ A comprehensive depletion map of the relevant TXTL resources in CFPS over time and their effects on system activity might give valuable clues towards how reactions can be purposefully tailored. Beyond application, however, such models grant insight into the fundamental mechanisms governing CFPS.

The timing of optimal 3-PGA addition is highly sensitive to the initial reaction conditions. We observed multiple effects that distorted the presented results in Figs. 5E and F, primarily the initial concentration of DNA in the reaction. While we produced the presented results at 2 nM DNA, we also carried out the same assays at 5 nM. At this increased DNA concentration, spike-in of 3-PGA caused a prolonged lag-phase as well as decreased rate and yield, compared to the control reaction without added 3-PGA (Fig. S25). Analysing the rate-yield trajectory of a spike-in assay at 5 nM DNA showed that the inverse U-shape of the curve collapsed into a straight diagonal instead (Fig. S26). The nature of these effects remains unclear, although plausible explanations have been put forward elsewhere. ^6^

### Translation is the main source of batch-to-batch variability

The impact of extract complexity extends beyond termination characteristics. Many factors influence reaction behaviour, such as the energy solution formulation^7–10^ or the extract source strain.^8^ However, reactions derived from extracts prepared according to the same protocol can also differ between laboratories and operators, and even multiple batches of extract produced by the same operator will not necessarily lead to identical reaction behaviour.^64,65^ This batch-to-batch variability limits the applicability of CFPS for development of new tools, which may be mitigated by a better understanding of the mechanisms driving it.^6,62,65^ We therefore asked whether it would be possible to identify the main processes driving batch-to-batch variability. Our approach was to parameterise our model with data from different extract batches and identify parameter variations that most impact process outcomes (Fig. 6A).

We designed a series of experiments to quantify differences between batches across transcription, translation and maturation (Fig. 6B): an assay testing different template DNA concentrations, an assay testing different ribosome concentrations, and an assay where we added translation inhibitor chloramphenicol to a reaction after 40 min runtime. We expected that the titrations of DNA and ribosomes would decouple parameters related to transcription and translation, respectively. Since chloramphenicol stalls elongating ribosomes, preventing production of new proteins, we used it to decouple translation from protein maturation. We conducted each experimental assay for all batches on the same day using the exact same reagents except for the extract solution, to ensure comparability, and combined the data from all assays per batch before feeding them into a Bayesian inference pipeline (see SI).

The fitted parameters were the initial energy concentration, *e*_0_, transcription and translation rate constants, *k_tx_* and *k_tl_*, mRNA and energy degradation rate constants, *k_d_* and *k_de_*, the maturation rate constant, *k_m_*, Michaelis-Menten coefficients for energy binding during transcription and translation, *K_tx_* and *K_tl_*, and transcription and translation energy consumption scalars, *q_x_* and *q_l_*.

Our baseline model with implemented energy degradation, parameterised from the posterior parameter distributions, recovers the main trends in the experimental data from each batch (Fig. 6B). The monotonic increase in activity from low to high concentrations of DNA and ribosomes as well as earlier termination at lower yield upon addition of chloramphenicol are generally captured in all batches. We quantified inter-batch variation for each parameter using the ratio 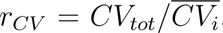, where *CV_tot_* is the the coefficient of variation *CV* = *σ/µ* over all three batches, and 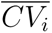 the mean of the coefficient of variation for individual batches (Fig. 6C). A large *r_CV_* denotes large inter-batch variation relative to per-batch variation. As such, large *r_CV_* indicates a parameter that has substantially shifted between batches. A small *r_CV_*, in turn, denotes small inter-batch variation or large per-batch variation, indicating a less substantial difference across batches.

**Figure 6:**
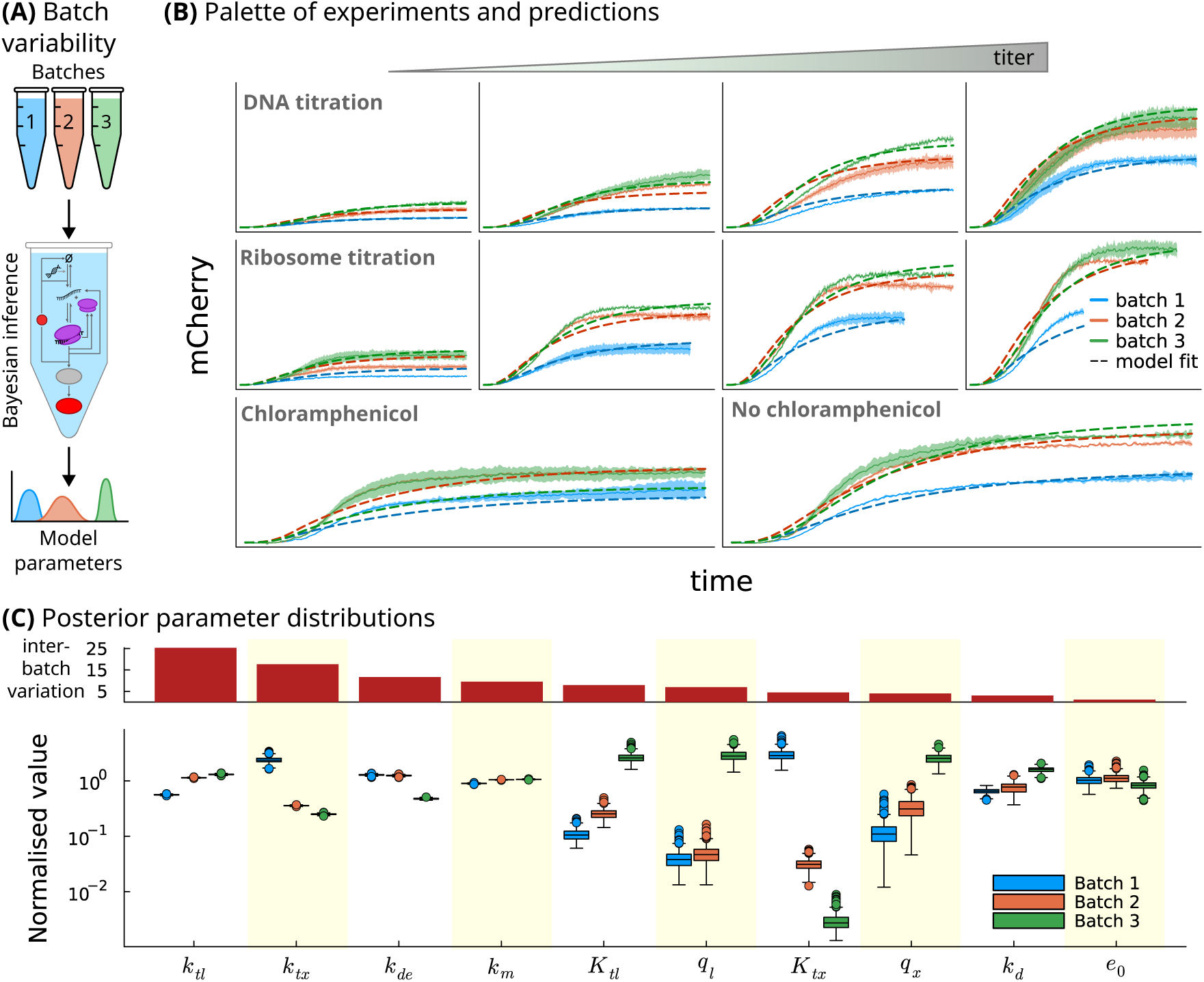
Batch-to-batch variability analysis. **(A)** Parameterisation of the baseline model with energy degradation on data from different batches of extract by Bayesian inference should yield insight into what processes, and therefore model parameters, differ between the batches. **(B)** We carried out three experimental assays (rows) for three batches of S150 extract (colours). Reactions ran at 0.25-2 nM template DNA (top row), 0.12-0.81 µM ribosomes (middle row) and with or without addition of chloramphenicol after 40 min (bottom row). Plotted are mCherry fluorescence over time (light colour, solid lines, shaded area is one standard deviation over triplicates) alongside predictions of the model with parameter values randomly sampled from the posterior parameter distributions after Bayesian inference (dark colour, dashed lines). Some experimental time series are shorter than others, which is considered by model predictions. **(C)** The posterior distributions of model parameters after Bayesian inference for batches 1-3 as boxplots. The three posterior distributions are normalised by their total mean for each parameter for better visualisation of trends. The inter-batch variation, as the ratio of total and mean of coefficients of variation, 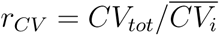, is shown as a barplot above.

The transcription and translation rate constants displayed the greatest inter-batch variation. The posteriors of the two parameters mirror each other: *k_tx_* decreases in direction of batch 1-3, whereas *k_tl_* increases. Although the parameter posteriors are not negatively correlated within each batch (Figs. S17a-S17c), a clear negative correlation is observed between batches (Fig. S17d). Because transcription and translation have redundant effects if only protein output is observed, their mirrored posteriors suggest incompletely decoupled parameters. The same trends are found in the Michaelis-Menten coefficients for transcription and translation, *K_tx_* and *K_tl_*, for similar reasons and because these are positively correlated with their respective rate constants (Fig. S17). The energy degradation rate constant *k_de_* and protein maturation rate constant *k_m_* display very tight posteriors which change slightly in their mean, resulting in a moderate *r_CV_* . The posterior distribution means of parameters related to energy affinity, *K_tl_* and *K_tx_*, and consumption, *q_l_* and *q_x_*, vary over orders of magnitude, but because their per-batch variations are also high, the overall inter-batch variation is low. Computational estimates for the initial energy concentration *e*_0_ show lowest inter-batch variation. This is consistent with experimental design, as most chemical energy was supplied in the form of energy solution, which was identical among the batches. In summary, the translation rate constant *k_tl_* shows the highest inter-batch variability, identifying translation as the process that varies the most between batches, according to the model parameterisation.

We made two further observations on the experimental and computational data. First, the model predictions for protein maturation seem to qualitatively deviate from the experimental observations. The model predicts signal to increase faster but terminate slower than is observed in the experimental data (Fig. 6B). This suggests a trade-off between capturing lag and termination phases using one maturation rate: A faster rate will lead to more abrupt termination but at the cost of not capturing the initial lag phase. The posterior distribution of the maturation rate *k_m_* is fairly constant throughout the batches (Fig. S16) but an order of magnitude lower than would have been expected from preliminary experiments tailored to measuring the maturation rate (see SI, Figs. S4, S16). In fact, *k_m_* is the only parameter with a posterior distribution that lies well outside the range of its prior (Fig. S16). The data therefore suggest that production of mature mCherry requires at least two processes for its description: a delay time, accounting for the lag phase at the beginning of the reaction, followed by a continuous maturation rate, capturing the gradual saturation of signal after expression termination. A possible explanation for these dynamics could be the formation of an intermediate to the mature fluorescent protein, as has been investigated in other reporters previously.^66^ Our baseline model, assuming only a continuous first-order rate, does not fully recover this (see SI).

Second, there appears to be an increase in activity from batch 1 through to batch 3 (Fig. 6B). Calculated as the average ratio between yields of each experiment, batch 2 has around 82 %, and batch 1 only 56 % of the activity of batch 3. The reason for this variation in activity is unknown, but different extract age and slight differences in extract preparation and reaction setup are viable causes, as previously studied.^67,68^ We note that the inferred distributions per parameter among batches shift monotonically with batch activity, either up or down. As determined above, we found the main change in inter-batch variation of the translation rate constant. However, the monotonic changes in all parameters indicate that, instead of identifying distinct parameter patterns for each batch, the model seemingly fits the same pattern to the respective batch activity globally. This should be considered when interpreting the presented results, as the resulting parameter values may not reflect the underlying process kinetics accurately.

## Discussion

In this work, we developed a framework for studying CFPS mechanisms using model-guided experimental design and used it to investigate various aspects of *in vitro* gene expression. We selectively expanded a coarse-grained baseline model of CFPS to answer isolated questions about different mechanisms underlying gene expression. Information gained from the isolated studies could be used to inform downstream experimental design decisions or be included in the model to increase its predictive power. The presented framework therefore lends itself well to DBTL-style system development.

We studied the rate-limiting steps in transcription and translation to find that the transcription rate seemed limited by both initiation and elongation, depending on the length of the gene, while elongation dominated the translation rate. A process that is rate-limited by elongation can be sped up greatly only by increasing the elongation rate,^69^ meaning that identification of the bottleneck can aid in optimising system activity. While we could mostly decouple initiation and elongation rates, the experimental data remains a product of more potential influences. As such, we attempted to minimise secondary effects on elongation rate, such as changes in sequence composition, by exactly copying and removing sequences, instead of introducing new ones. Distortion of the translation rate, e.g. by truncation, is a known effect and warrants further analysis towards the exact mechanism and locations of truncation. Alternatively, the same analysis may be performed via a different and more direct method of protein monitoring, such as using a shorter luciferase reporter,^70^ a halo-tag^71^ or use of fluorescent lysine-BODIPY readouts.^72^ We assumed T7 RNAP drop-off akin to protein truncation to be negligible here, due to a lack of published evidence to the contrary. However this, too, may be investigated, e.g. by use of dual aptamers at the 5’ and 3’ end of the transcript, as drop-off is an alternative explanation for the length dependence of the transcription rate. It would be interesting to further decouple the contributions of initiation and termination, instead of treating them as one lumped rate, by selectively altering only the promoter or terminator sequence and observing the effect on overall rate.^41^ Further follow-up questions are the dependence of these results on the system used, e.g. by using a different polymerase. The characterisation of these phenomena would further advance our understanding of transcription and translation rates. Finally, we encountered an issue when measuring mRNA levels over time, which we suspect to relate to the degradation of the DFHBI-1T dye as the reaction proceeds. However, the early timepoints used for calculation of the slope are not affected by this (Fig. S27).

Next, we investigated termination of batch CFPS reactions. In a first step, we identified reaction lifetime to be mainly limited by a process unrelated to TXTL activity, which later turned out to be connected to the degradation of the chemical energy carriers, NTPs, and their secondary energy source 3-PGA. Again, the presented results seem highly dependent on the system used. It is unclear, why the system studied here is terminated by depletion of chemical energy whereas previous studies of extract-based systems found other limiting resources, such as amino acids.^7,58–61^ It is likely that both the extract as well as the energy systems play a role in determining the first limiting substrate and further deconvolution of this network may give important insight into how reactions can be run most efficiently. The complexity of extract CFPS makes such predictions unfeasible *a priori*, requiring instead experimental characterisation. Fully defined systems such as PURE represent a more promising platform for thorough predictability of resource depletion.^15,36^

Finally, we parameterised the model on three batches of extract via a limited set of reactions. We identified the translation rate constant *k_tl_* as the parameter responsible for the largest change in model predictions; however mechanistic insight into this trend is limited. The translation rate may vary due to differences in enzyme concentration, chemical environment, or due to a mechanistically entirely different effect, which the model can best fit by changing this parameter. Whether it is indeed the translation rate in the *in vitro* reactions that changes between batches cannot be conclusively asserted. Since the model is coarse-grained, any mechanism that goes beyond its scope cannot be elucidated. We observed this with the maturation rate *k_m_*, which had to fit a phenomenon potentially linked to two mechanisms and as a result was not able to fit either well (see SI, also considered previously^73^). Global changes, potentially caused by e.g. deterioration over storage time, seemed to be dominant over more subtle differences between extract batches. The model captured this trend by monotonically varying all parameters to adapt the protein synthesis activity. However, whether all of them also change correspondingly in the reactions, is doubtful. Indeed, remaining parameter correlation, alongside the opposing trends of transcription and translation rate constants, showed that the performed experiments were not sufficient to fully decouple the coarse-grained model parameters. It therefore seems necessary to carry out more sophisticated experimental investigations to dissect the parameter space and identify more subtle batch differences.^37,74^ This might be achieved by ensuring decoupling of parameters during the experimental design process using sensitivity analysis,^69,75,76^ as well as measuring more outputs, like RNA, to further constrain the model.

Overall, we demonstrated that targeted analysis using tailored alterations of a baseline model works well to test specific hypotheses, while untargeted analysis to find *a priori* unknown patterns in a limited dataset by inference on the baseline model leaves much room for interpretation. The use of models to guide experimental design of CFPS work is established in the field.^15,16,18,35,36,69^ For complete control over reaction conditions and model-guided design, reconstituted systems such as PURE may be better suited than crude ones.^15,16,36,77^ While the variable and noisy nature of extract-based CFPS means that it will likely never be fully characterisable, models can help to alleviate this shortcoming and improve the usability of CFPS as a tool to molecular biologists.^6,14,74^

## Methods

### S150 extract preparation

We mostly adopted the extract preparation protocol from^3^ with slight modifications. BL21(DE3) star lacZ::frt wtASD rRNA, kindly supplied by,^3^ were used to inoculate 10 mL 2x YTPG (Formedium, TRP01; Millipore, 1.03753.0500; Fisher Scientific, S/3160/63; Sigma-Aldrich, P5379-500G; Sigma-Aldrich, P3786-500G) with Kanamycin (Sigma-Aldrich, K1377) and incubated at 37 *^◦^*C, 220 rpm overnight. The culture was used to inoculate 4x 400 mL 2x YTPG with Kanamycin and incubated at 37 *^◦^*C, 220 rpm until OD_600_ 0.5. The culture was induced to 0.1 mM IPTG (Thermo Scientific, R0392) and incubation was continued until OD_600_ 3. The culture was cooled down on ice and centrifuged at 5000 g, 4 *^◦^*C for 15 min. The supernatant was decanted and the cells washed once in S150A [20 mM Tris, 100 mM NH4Cl, 10 mM MgCl2, 0.5 mM EDTA, 2 mM DTT, pH 7.2], before being pelleted at 6000 g, 4 *^◦^*C for 10 min and flash-frozen in liquid nitrogen. The frozen pellet was thawed and resuspended in 10 mL S150A at 10 % (w/v). Protease Inhibitor (Thermo Scientific, 78425) was added to 4.5 µL g*^−^*^1^. The cells were disrupted at 20000 psi in a cell disruptor (Constant Systems, CF-1). The lysate was loaded on top of 5 mL S150A and centrifuged at 30000 g, 4 *^◦^*C for 30 min in a Beckman XPN-100 ultracentrifuge using 14 mL tubes (Seton, 5031) in a SW40 rotor (Beckman). The process was repeated with the upper two thirds of supernatant at 90000 g for 21 h and again at 141000 g for 3 h. The upper two thirds of the final supernatant were dialysed against 1 L S150B [50 mM HEPES, 10 mM Mg-Glu, 200 mM K-Glu, 2 mM DTT, 1 mM spermidine, 1 mM putrescine, pH 7.6] using a 3.5K MWCO Snakeskin membrane (XXX) for 3x 2 h and 1x overnight, exchanging the dialysate in between. The dialysed solution was concentrated down to 90-100 mg mL*^−^*^1^, measured by Nanodrop in BSA mode, in 3K Amicon Ultra-15 (EMD, UFC900324). The resulting S150 extract was aliquoted in appropriate volumes, flash-frozen in liquid nitrogen and stored at -80 *^◦^*C until usage.

### Cloning and plasmid preparation

All plasmids used for this study were derived from pEXP5.^35^ pCW004 (dmCherry-2xdBroccoli) was cloned by AQUA^78^ from pEXP5 through a M71E mutation.^50^ pCW005 (dmCherry-mCherry-2xdBroccoli) was cloned by AQUA using pEXP5 as backbone and dmCherry from pCW004 as insert. pCW006 (dmCherry-dmCherry-mCherry-2xdBroccoli) was cloned by Golden Gate, using pCW005 as backbone and dmCherry from pCW004 as insert. pCW009 (halfmCherry-2xdBroccoli), pCW010 (2xdBroccoli) and pCW011(3xRBS-2xdBroccoli) were all cloned by AQUA from pEXP5. All plasmids were amplified by Maxiprep (Zymo, D4202) for maximal purity and diluted down to 25 nM with water.

### CFPS reactions

Reactions were combined from energy solution, S150 extract, 70S ribosomes, DNA, and aptamer dye. A 4x energy solution was prepared,^79^ with Mg-Glu at 10 mM, K-Glu at 100 mM, and tRNAs omitted. S150 extract was prepared as above and used at a final reaction concentration of 10 mg mL*^−^*^1^. 70S ribosomes were prepared according to^80^ and used at a final reaction concentration of 500 nM. DNA was prepared as above and used at a final reaction concentration of 2 nM. DFHBI-1T (Tocris, 5610) was dissolved in DMSO and diluted to a working stock concentration of 1 mM with water. The reaction concentration was 100 µM. Unused energy solution, extract and ribosomes were freeze-thawed a maximum of three times, otherwise they were discarded. Reactions were carried out in technical triplicates at 5 µL volume in 384 well plates (Greiner, 784076), sealed with plate film (Excel Scientific, STR-SEAL-PLT), at 37 *^◦^*C in a platereader (Agilent, Biotek Synergy H1 or BMG Labtech, Fluostar Omega). Reactions were measured at 485/520 nm for Broccoli signal and 580/620 nm for mCherry signal every 2 min over 12 h with 300 rpm shaking in between readings. On the Fluostar, 1000 gain was used for both fluorophores. On the Biotek, Broccoli was measured at gain 50 and mCherry at gain 70.

### Data processing

Raw data in RFU was transformed into micromolar units by calibrating curves for the aptamer and the protein (Fig. S18). Where necessary, due to dilution of the reaction volume by spike-in, micromolar units were further transformed into mass units. Broccoli data required background subtraction due to strong autofluorescence from the extract. A negative control without DNA template was used for background subtraction. Linear fits were calculated via a manually implemented weighted least-squares script.

### Model simulations

The model was implemented in Julia^81^ as a system of ordinary differential equations and solved with DifferentialEquations.jl^82^ using the Rodas5 solver and the state saved every 2 min, emulating the platereader measurements. Spike-in experiments were simulated via callback functions, diluting all species and increasing the concentration of the spiked-in species.

### Bayesian inference

Bayesian parameter inferences was performed using Turing.jl.^83^ Prior parameter distributions were estimated by literature values (Tab. S1). Parameters were assumed normally distributed unless *µ <* 6*σ*, in which case a Lognormal distribution was used to ensure positivity. During inference, all parameter distributions were scaled to 1, and scaled back up for predictions. The likelihood was calculated internally in Turing.jl as the logarithm of the probability density function of a Normal distribution with the model prediction at any time point as mean and the experimental uncertainty as standard deviation, evaluated at the observed experimental data. The log-likelihood was then summed up over all time points. Data between experimental series was normalised to common reference experiments, to minimise the influence of batch effects. The Turing.jl implementation of the MCMC ensemble sampler emcee^84^ was used with 100 walkers and stretch length 2. Inference was run until confirmed to be converged visually and by use of the Geweke diagnostic and autocorrelation time. Walkers three standard deviations outside the mean of the final distribution were excluded from further analysis to prevent bias from non-converged walkers. The final 100 steps of the remaining walkers were used to estimate the posterior parameter distribution.

### Gaussian Process Regression

Gaussian Process Regression was done using AbstractGPs.jl, separately for the rate and yield dimensions. A squared exponential kernel with a length scale of 100 min was applied on each dimension and resulting predictions were plotted against each other to produce the two-dimensional trajectory. Uncertainties were calculated as ellipses drawn with the standard deviation in both dimensions at each point of the prediction.

## Acknowledgement

The authors thank A. Pompa for assistance in using the ultracentrifuge, which was supported by funding from the Wellcome Discovery Research Platform for Hidden Cell Biology [226791], and the Edinburgh Protein Production Facility for access to a cell disruptor, both used in the preparation of cellular extract. Detailed protocols for preparation of S150 extract were provided by Prof. Wataru Aoki. Preparation of purified ribosomes was aided by Dr. Sahan Liyanagedera. We thank Prof. Dora Tang for providing the pEXP5 DNA construct. The cloning of subsequent DNA templates was assisted by Dr. Alexandru Popov. Finally, we thank Rodrigo Lara Molina for insightful discussions on Bayesian inference and Elena Pascual Garcia for advice on continuous modelling of polysomes.

## Supporting Information Available

### Baseline model

The full formulation of the baseline model was

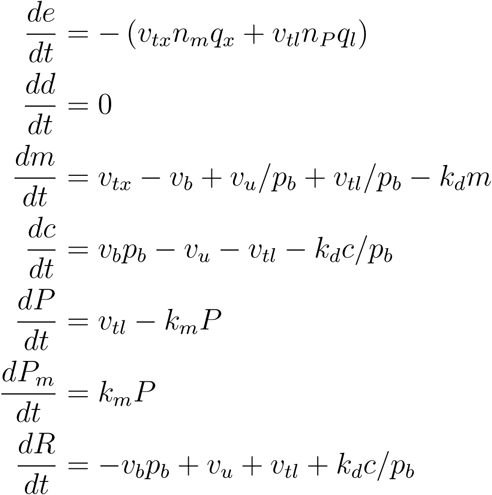

with rates

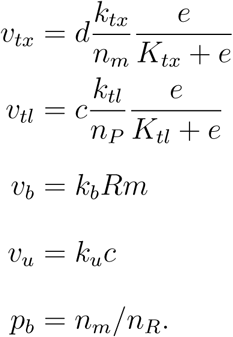

We determined the rate constants from literature values or manually extracted from experiments and estimated initial concentrations of model species from the experimental setup. Both are listed in Tab. S1.

### T7 RNAP abundance

To our knowledge, T7 RNAP polymerase abundance in extract induced with IPTG to produce the enzyme has not been measured before. However, an idea of what enzyme concentration is to be expected would be valuable for modelling purposes. Therefore, we roughly estimated the number based on literature values of T7 RNAP yields after harvesting the protein from an induced culture by His-tag purification. A few papers have reported values of T7 RNAP yield in units of grams.^85–88^ Knowing their respective culture densities and comparing them to the typical density of the extract prepared for this work, we derived an estimate for T7 RNAP in the used S150 extract on the order of 10 µM (Tab. S2). We calculated this assuming 8 *·* 10^8^ *E. coli* cells per millilitre of culture at OD_600_, a wet weight of 1 *·* 10*^−^*^12^ per cell, and a molecular weight of T7 RNAP of 99 000 g mol*^−^*^1^. Since we used a DNA concentration of a few nanomolar in the reactions, we therefore considered T7 RNAP abundant and not necessary to model explicitly.

**Table S1:**
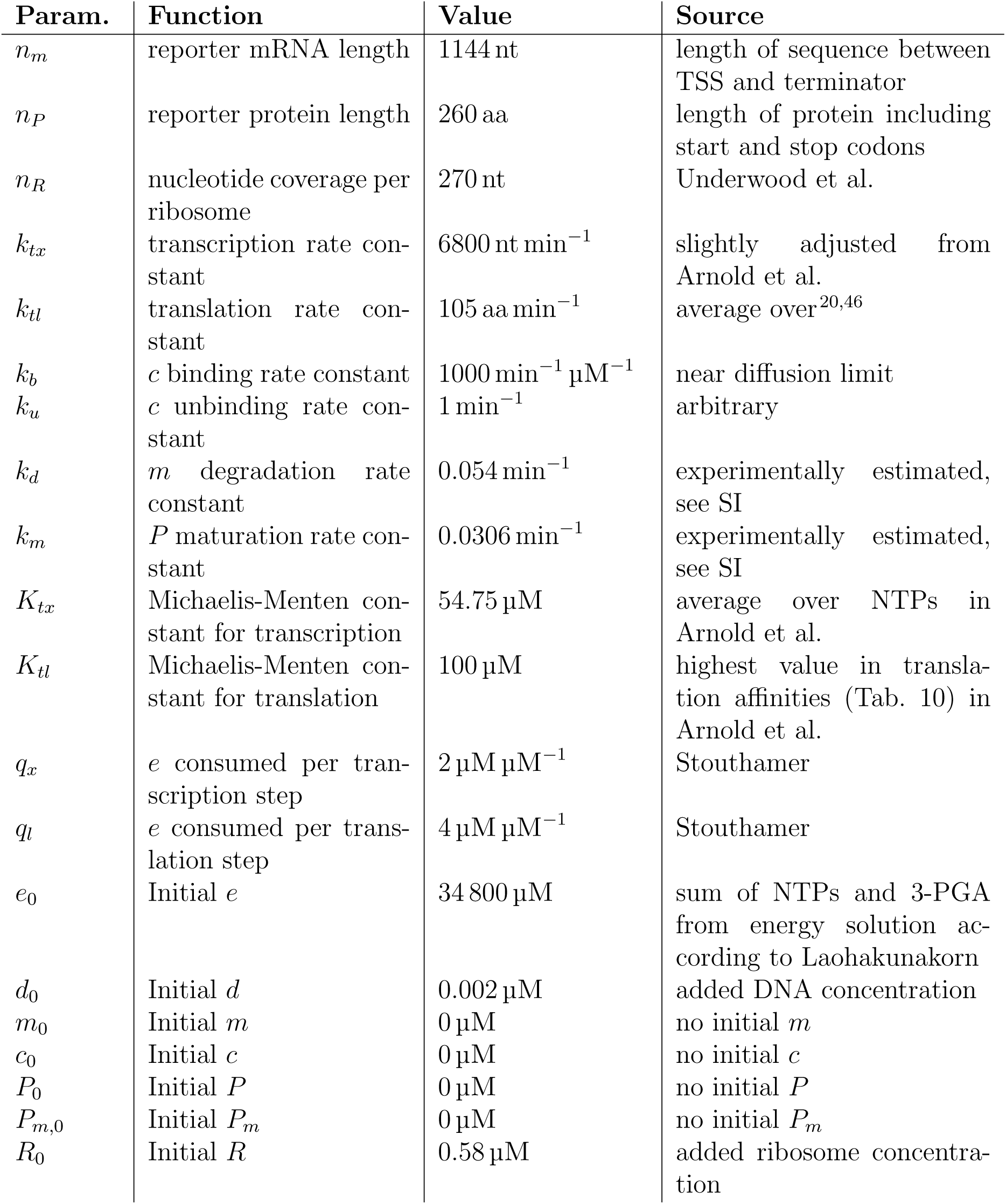
Default model parameters and their sources.

| **Param.** | **Function** | **Value** | **Source** |
|---|---|---|---|
| *n*<sub>*m*</sub> | reporter mRNA length | 1144 nt | length of sequence between TSS and terminator |
| *n*<sub>*P*</sub> | reporter protein length | 260 aa | length of protein including start and stop codons |
| *n*<sub>*R*</sub> | nucleotide coverage per ribosome | 270 nt | Underwood et al. |
| *k*<sub>*tx*</sub> | transcription rate con-stant | 6800 nt min<sup>−1</sup> | slightly adjusted from Arnold et al. |
| *k*<sub>*tl*</sub> | translation rate con-stant | 105 aa min<sup>−1</sup> | average over<sup>20,46</sup> |
| *k*<sub>*b*</sub> | *c* binding rate constant | 1000 min<sup>−1</sup> µM<sup>−1</sup> | near diffusion limit |
| *k*<sub>*u*</sub> | *c* unbinding rate con-stant | 1 min<sup>−1</sup> | arbitrary |
| *k*<sub>*d*</sub> | *m* degradation rate constant | 0.054 min<sup>−1</sup> | experimentally estimated, see SI |
| *k*<sub>*m*</sub> | *P* maturation rate con-stant | 0.0306 min<sup>−1</sup> | experimentally estimated, see SI |
| *K*<sub>*tx*</sub> | Michaelis-Menten con-stant for transcription | 54.75 µM | average over NTPs in Arnold et al. |
| *K*<sub>*tl*</sub> | Michaelis-Menten con-stant for translation | 100 µM | highest value in transla-tion affinities (Tab. 10) in Arnold et al. |
| *q*<sub>*x*</sub> | *e* consumed per tran-scription step | 2 µM µM<sup>−1</sup> | Stouthamer |
| *q*<sub>*l*</sub> | *e* consumed per trans-lation step | 4 µM µM<sup>−1</sup> | Stouthamer |
| *e*<sub>0</sub> | Initial *e* | 34 800 µM | sum of NTPs and 3-PGA from energy solution ac-cording to Laohakunakorn |
| *d*<sub>0</sub> | Initial *d* | 0.002 µM | added DNA concentration |
| *m*<sub>0</sub> | Initial *m* | 0 µM | no initial *m* |
| *c*<sub>0</sub> | Initial *c* | 0 µM | no initial *c* |
| *P*<sub>0</sub> | Initial *P* | 0 µM | no initial *P* |
| *P*<sub>*m*,0</sub> | Initial *P*<sub>*m*</sub> | 0 µM | no initial *P*<sub>*m*</sub> |
| *R*<sub>0</sub> | Initial *R* | 0.58 µM | added ribosome concentra-tion |

**Table S2:** T7 RNAP concentration as estimated from the literature.

| Reference | <sup>85</sup> | <sup>86</sup> | <sup>87</sup> | <sup>88</sup> |
|---|---|---|---|---|
| Culture wet weight (g) | 10 | 1 | 8 | 0.3 |
| Culture cell cout (cells) | 1 · 10<sup>13</sup> | 1 · 10<sup>12</sup> | 8 · 10<sup>12</sup> | 3 · 10<sup>11</sup> |
| Protein yield (g) | 0.06 | 0.015 | 0.05 | 0.00075 |
| Protein yield (mol) | 6 · 10<sup>−7</sup> | 1.5 · 10<sup>−7</sup> | 5 · 10<sup>−7</sup> | 7.5 · 10<sup>−9</sup> |
| Protein yield (count) | 3.6 · 10<sup>17</sup> | 9.1 · 10<sup>16</sup> | 3 · 10<sup>17</sup> | 4.6 · 10<sup>15</sup> |
| Count per cell | 36500 | 91250 | 38000 | 15200 |
| Count in extract | 1.4 · 10<sup>17</sup> | 3.5 · 10<sup>17</sup> | 1.5 · 10<sup>17</sup> | 5.6 · 10<sup>16</sup> |
| Concentration in extract (µM) | 232 | 581 | 242 | 97 |
| Concentration in reaction (µM) | 56 | 140 | 58 | 23 |

### mRNA degradation rate constant estimation

We estimated the rate constant of mRNA degradation, *k_d_*, experimentally as follows. We set up a CFPS reaction with purified mRNA instead of DNA in triplicate alongside two control reactions: a background control where we omitted DFHBI-1T and a folding dynamics control without extract (Fig. S1a). mRNA degrades in extract,^22^ however its effect is obscured by other dynamics, among them autofluorescence by the lysate and folding plus maturation of the aptamer. We therefore corrected the raw signal for autofluorescence and the folding dynamics (Fig. S1b). We then extracted a window of the resulting corrected signal between 20-60 min and calculated a weighted least squares fit on the data in log-scale (Fig. S1c). This gave a degradation rate constant of *k_d_* = 0.054 min*^−^*^1^ *±* 0.0035. Plotted in linear space, this recovers the data well (Fig. S1d).

We repeated this procedure for each batch in the batch-to-batch variability study (Fig. S2. Too little extract for S150_1_ was left to for the amount of reactions that we needed, leading to an apparently slower mRNA degradation rate, as we had added less extract than required. Therefore, we did not use the data in the Bayesian inference pipeline.

**Figure S1:**
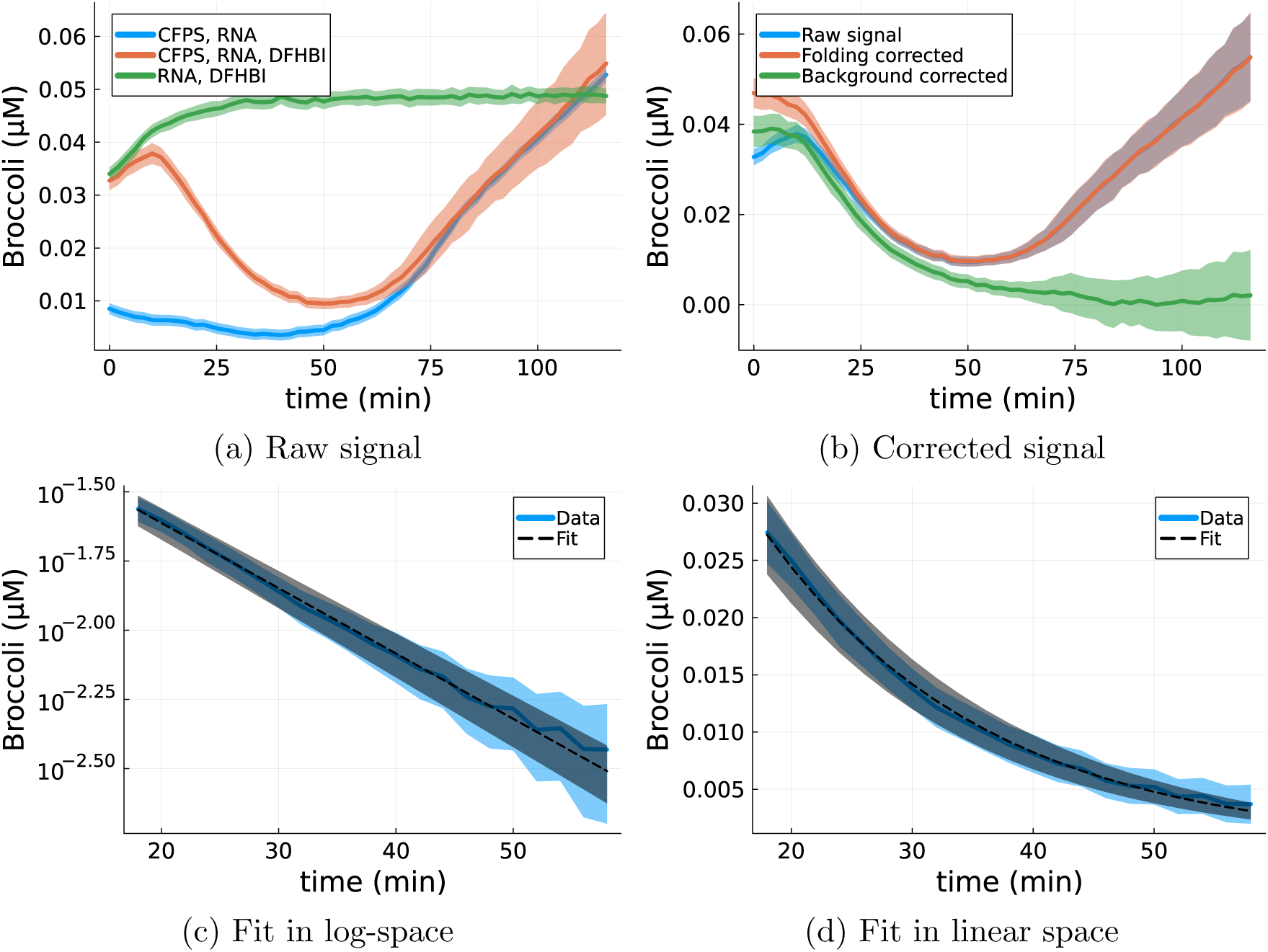
Experimental estimation of *k_d_*. (a) mRNA degradation signal (orange) obscured by background fluorescence (blue) and the aptamer maturation dynamics (green). Solid lines show average and shaded area one standard deviation over triplicates. (b) Obscured mRNA degradation (blue), after correction for folding dynamics (orange) and background-corrected (green). (c) Weighted least squares fit in log-space (dashed line) with standard deviation as grey shaded area and data in blue. (d) Same as (c) but in linear space.

**Figure S2:**
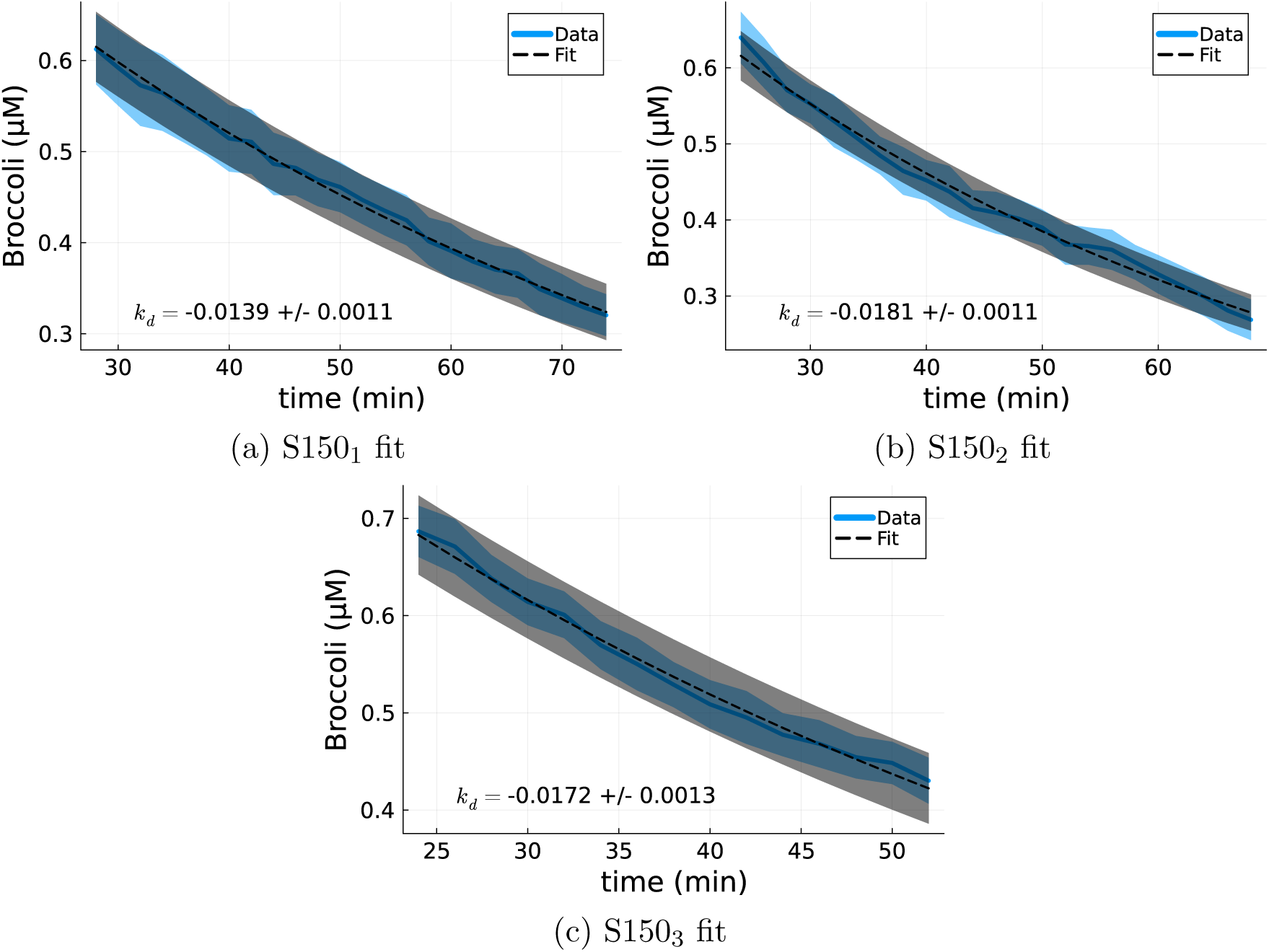
Weighted least squares fit of mRNA degradation experiments analogous to Fig. S1d, for S150_1_-S150_3_ (a-c). The reaction prepared with S150_1_ contained too little extract, wherefore the apparent degradation rate was lower. Solid lines show the average and shaded area one standard deviation over duplicates.

### Protein maturation rate constant estimation

Fluorescent proteins need time to mature before they can be observed, which models need to consider for accurate predictions.^66^ Maturation in CFPS is usually modelled as a first-order rate, dependent on a rate constant *k_m_*, which is multiplied on the pool of non-mature protein, to produce mature protein. ^14,16,35^ According to this approach, a population of non-mature proteins that are produced simultaneously will exponentially but gradually mature with a half-life depending on the rate constant (Fig. S3a).

**Figure S3:**
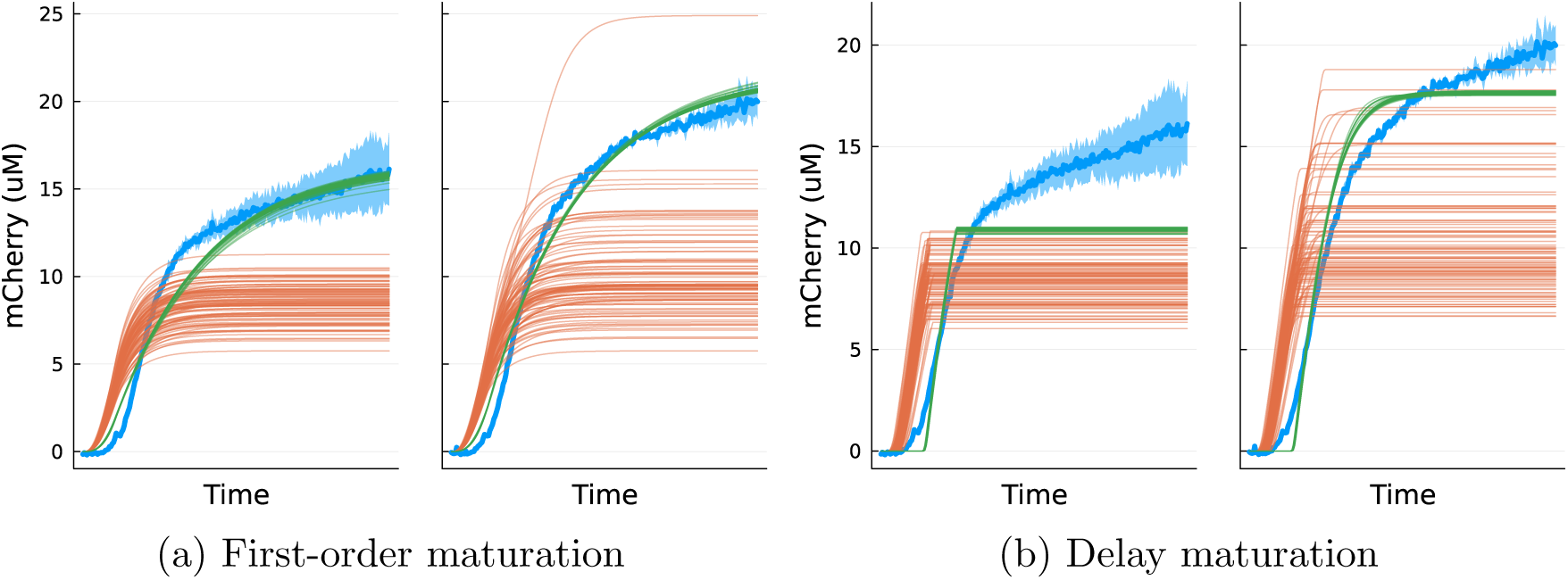
Inference of chloramphenicol addition experiment under competing maturation mechanism hypotheses. Data is shown in blue with the average as a solid line and its standard deviation as a shaded area over duplicates. Model predictions based on sampling from the parameter priors are shown in orange, while predictions based on samples from the posterior after Bayesian inference are shown in green. The left plot is data with chloramphenicol addition, the right plot a positive control with addition of water. (a) shows inference based on a first-order maturation rate while (b) is based on a discrete delay time.

An alternative approach is that of a discrete delay time *τ*, which specifies how long a non-mature protein takes, to mature.^18^ The maturation dynamics under this approach are qualitatively different from the first-order rate, as a pool of non-mature proteins that were produced simultaneously would mature at exactly the same time in the case of a discrete delay (Fig. S3b). Discrete delays can be implemented in the model as a delay differential equation.

We considered both approaches for this work. However, because the discrete delay by itself caused immediate termination *t* minutes after simulation of chloramphenicol addition, which we did not observe in the data (Fig. S3b), we chose the first-order maturation rate approach for the model.

To find a preliminary estimate for the maturation rate of mCherry, we took an approach similar to that of the mRNA degradation estimation. We prepared and measured six default reactions in the platereader. After 36 min, 1 µL we added chloramphenicol to three of the reactions at 100 µg mL*^−^*^1^, which has previously been shown to lead to inhibition of 80 % of ribosomes in CFPS.^89^ This should immediately stall elongating ribosomes, to prevent production of any new protein. Any observed increase in signal thereafter should be due to protein maturation exclusively. To the other three control reactions, we added 1 µL H_2_O. The signal showed an earlier termination in the sample with Chloramphenicol (Fig. S4a). We extracted the signal between 80-160 min, flipped it vertically and transformed it to log-space for a weighted least-squares fit, giving a maturation rate constant of *k_m_* = 0.0306 min*^−^*^1^ *±* 0.0027 (Fig. S4b). Converted back into linear space, the fitted curve recovers the data (Fig. S4c).

**Figure S4:**
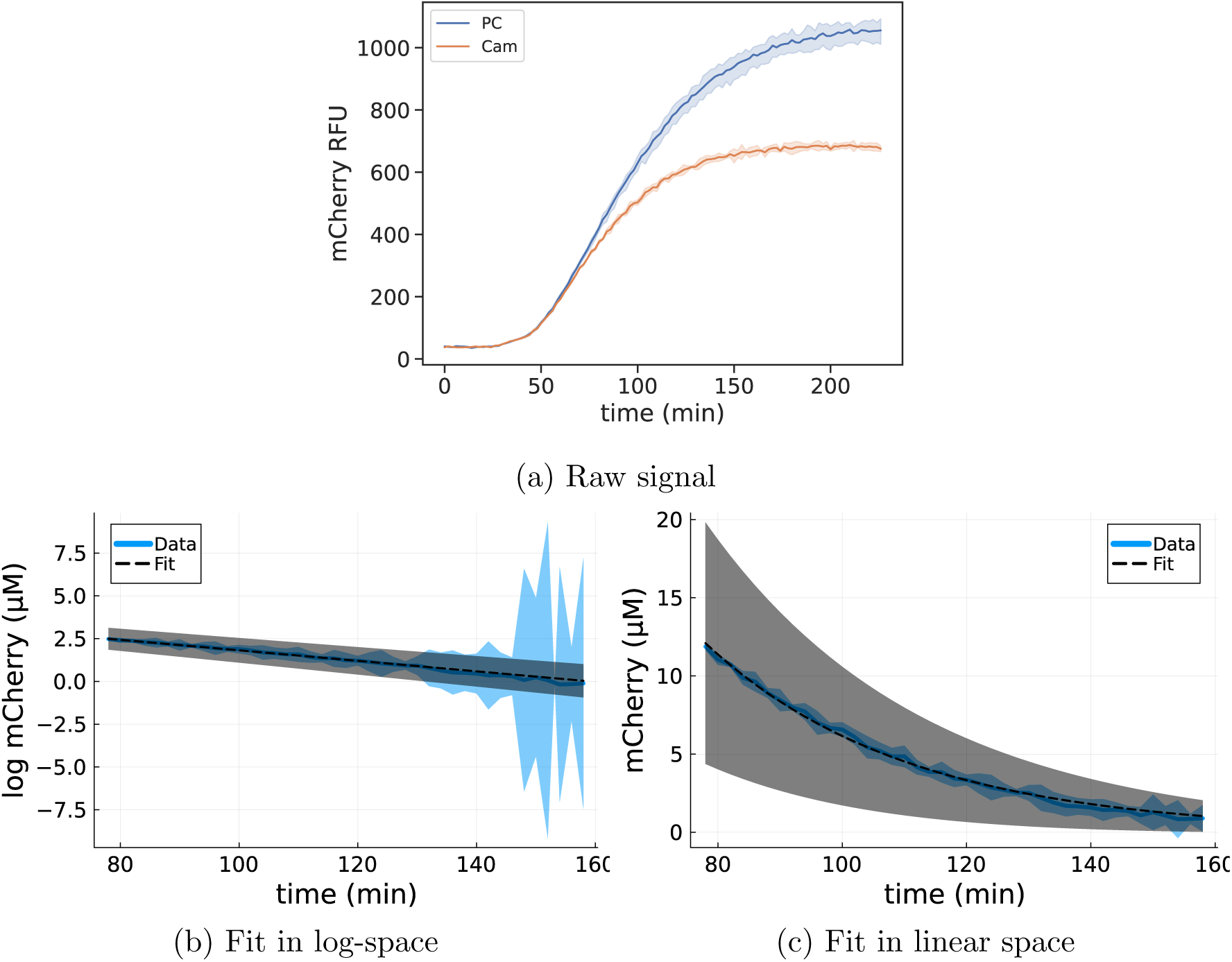
Experimental estimation of *k_m_*. (a) Reactions inhibited with chloramphenicol at 36 min (orange) terminated earlier than the default reactions without inhibition (blue). Solid lines shows the average and shaded area one standard deviation over triplicates. (b) The logarithm of the vertically flipped chloramphenicol data between 80-160 min (blue) with the fitted linear function (dashed line) and its standard deviation (grey shaded area) overlaid. (b) Same as in (b) but in linear space.

## Bayesian inference

### Rate limitation inference

We carried out inference as described in the methods. We expanded the model for an mRNA maturation step analogous to protein maturation, set to 0.06 min*^−^*^1^,^90^ and extracted the transcription rate from simulations by finding the highest rate of change of the sum of mature free mRNA and mature mRNA in complex with the ribosome. Because the concentration of mRNA is on a much smaller scale than the parameters to be inferred, we scaled up the data and model predictions by a factor of 10,000. The protein data was scaled up by a factor of 100 to achieve the same.

**Table S3:**
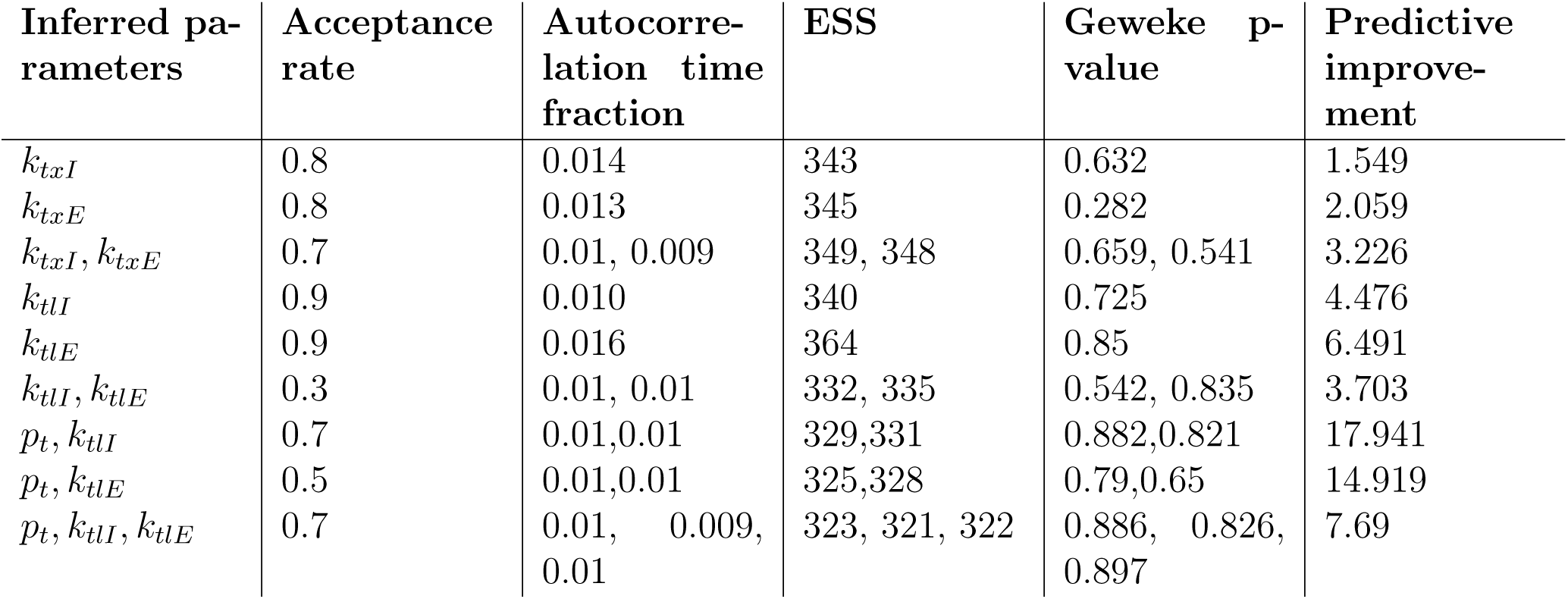
Convergence metrics for rate limitation study.

| **Inferred pa-rameters** | **Acceptance rate** | **Autocorre-lation time fraction** | **ESS** | **Geweke p-value** | **Predictive improve-ment** |
|---|---|---|---|---|---|
| *k*<sub>*txI*</sub> | 0.8 | 0.014 | 343 | 0.632 | 1.549 |
| *k*<sub>*txE*</sub> | 0.8 | 0.013 | 345 | 0.282 | 2.059 |
| *k*<sub>*txI*</sub>, *k*<sub>*txE*</sub> | 0.7 | 0.01, 0.009 | 349, 348 | 0.659, 0.541 | 3.226 |
| *k*<sub>*tlI*</sub> | 0.9 | 0.010 | 340 | 0.725 | 4.476 |
| *k*<sub>*tlE*</sub> | 0.9 | 0.016 | 364 | 0.85 | 6.491 |
| *k*<sub>*tlI*</sub>, *k*<sub>*tlE*</sub> | 0.3 | 0.01, 0.01 | 332, 335 | 0.542, 0.835 | 3.703 |
| *p*<sub>*t*</sub>, *k*<sub>*tlI*</sub> | 0.7 | 0.01,0.01 | 329,331 | 0.882,0.821 | 17.941 |
| *p*<sub>*t*</sub>, *k*<sub>*tlE*</sub> | 0.5 | 0.01,0.01 | 325,328 | 0.79,0.65 | 14.919 |
| *p*<sub>*t*</sub>, *k*<sub>*tlI*</sub>, *k*<sub>*tlE*</sub> | 0.7 | 0.01, 0.009, 0.01 | 323, 321, 322 | 0.886, 0.826, 0.897 | 7.69 |

**Figure S5:**
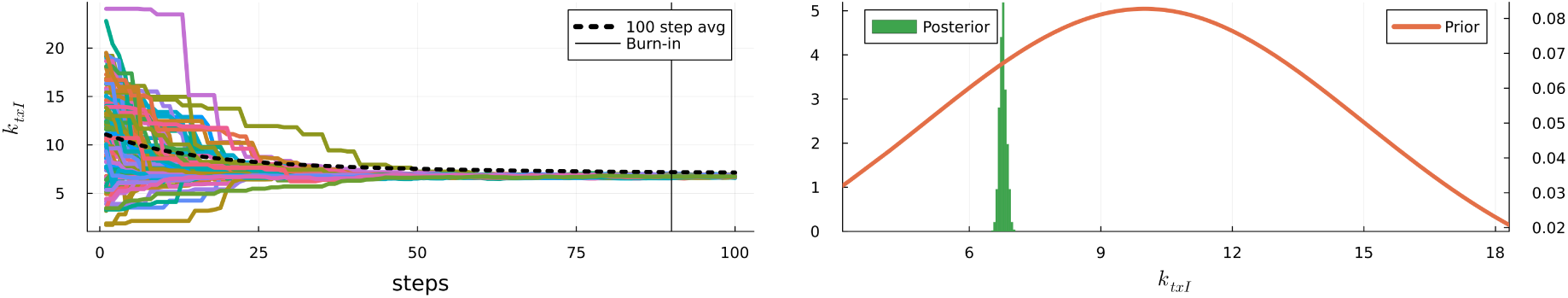
Raw chain trajectories and posterior for *k_txI_*, assuming an initiation-limited regime. (left) Raw walkers (coloured lines), average over all walkers over the last 100 steps (dashed line), and burn-in steps marked with a vertical line. (right) Density function of the prior distribution (orange, right y-axis), compared to posterior distribution after burn-in (green, left y-axis).

**Figure S6:**
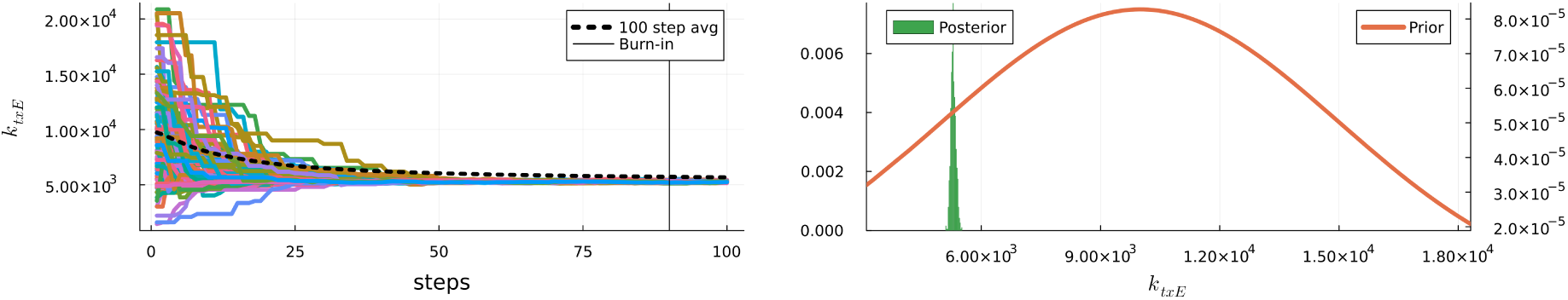
Raw chain trajectories and posterior for *k_txE_*, assuming an elongation-limited regime. (left) Raw walkers (coloured lines), average over all walkers over the last 100 steps (dashed line), and burn-in steps marked with a vertical line. (right) Density function of the prior distribution (orange, right y-axis), compared to posterior distribution after burn-in (green, left y-axis).

**Figure S7:**
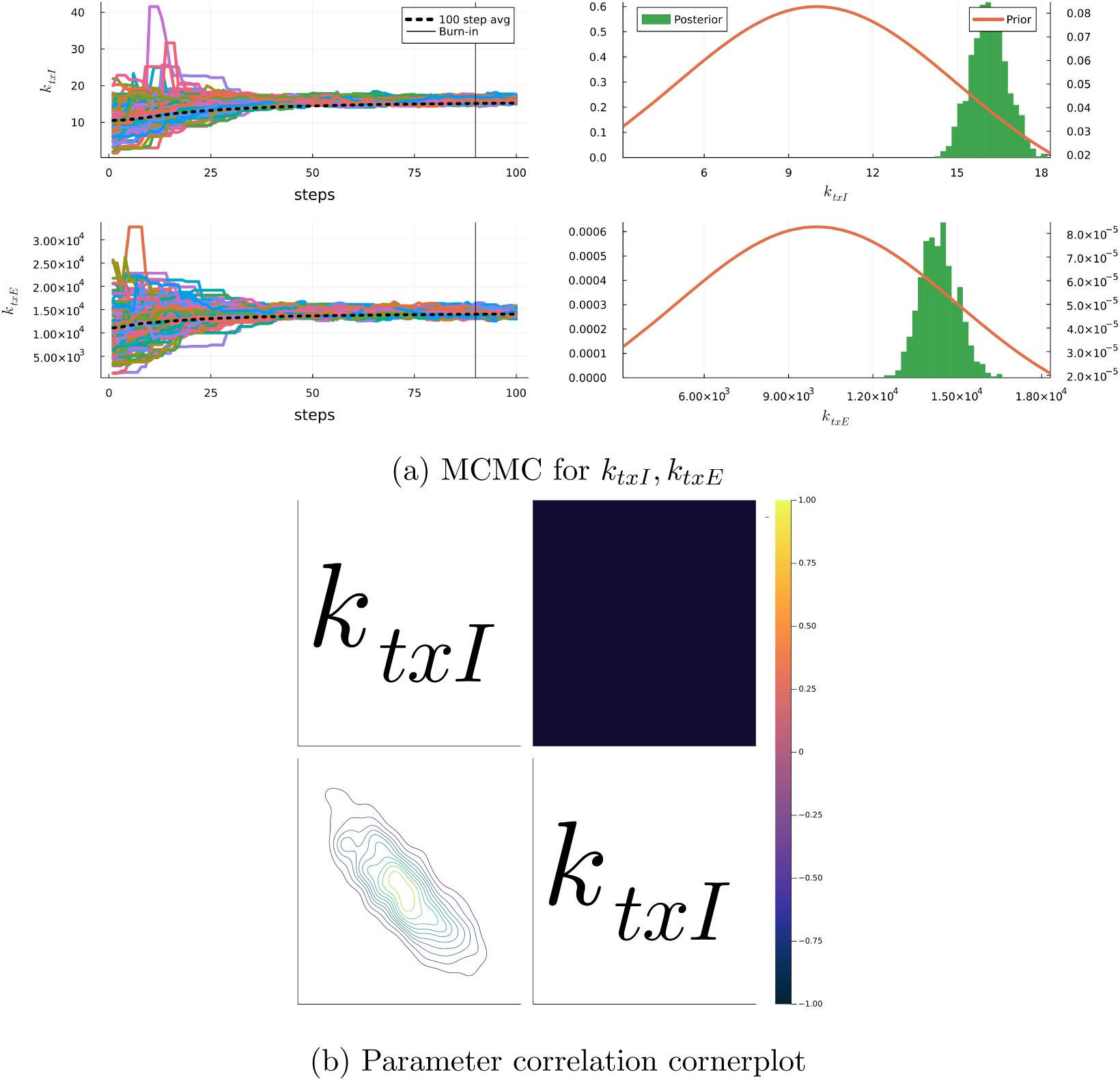
(a) Raw chain trajectories and posterior for *k_txI_, k_txE_*, assuming contributions from both. (left) Raw walkers (coloured lines), average over all walkers over the last 100 steps (dashed line), and burn-in steps marked with a vertical line. (right) Density function of the prior distribution (orange, right y-axis), compared to posterior distribution after burn-in (green, left y-axis). (b) Cornerplot of posterior correlations. Contour plot showing posterior correlations (lower left). Correlation in linear-linear (upper left triangle) and log-log space (lower right triangle) (upper right), according to colourbar on the right.

**Figure S8:**
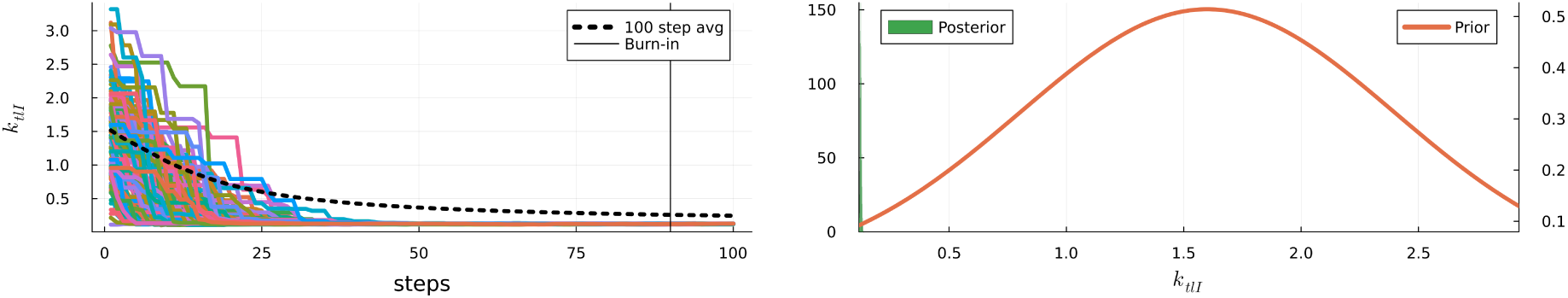
Raw chain trajectories and posterior for *k_tlI_*, assuming an initiation-limited regime. (left) Raw walkers (coloured lines), average over all walkers over the last 100 steps (dashed line), and burn-in steps marked with a vertical line. (right) Density function of the prior distribution (orange, right y-axis), compared to posterior distribution after burn-in (green, left y-axis).

**Figure S9:**
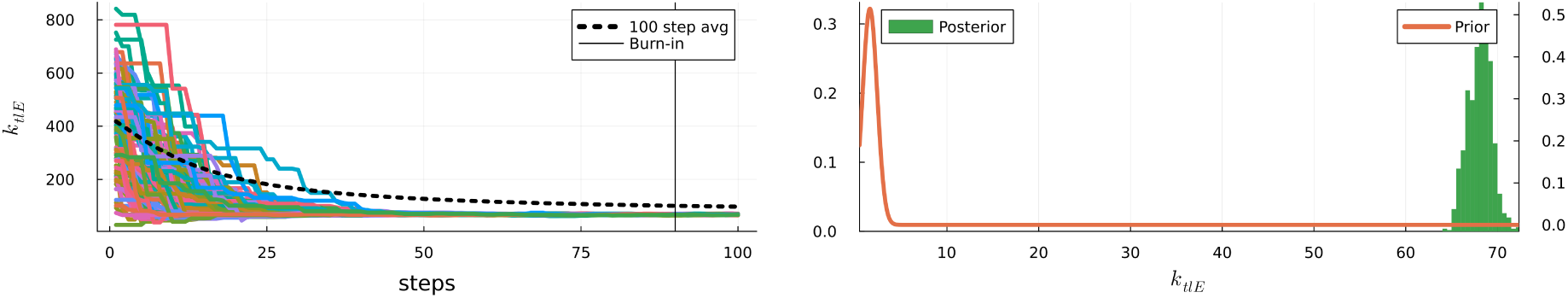
Raw chain trajectories and posterior for *k_tlE_*, assuming an elongation-limited regime. (left) Raw walkers (coloured lines), average over all walkers over the last 100 steps (dashed line), and burn-in steps marked with a vertical line. (right) Density function of the prior distribution (orange, right y-axis), compared to posterior distribution after burn-in (green, left y-axis).

**Figure S10:**
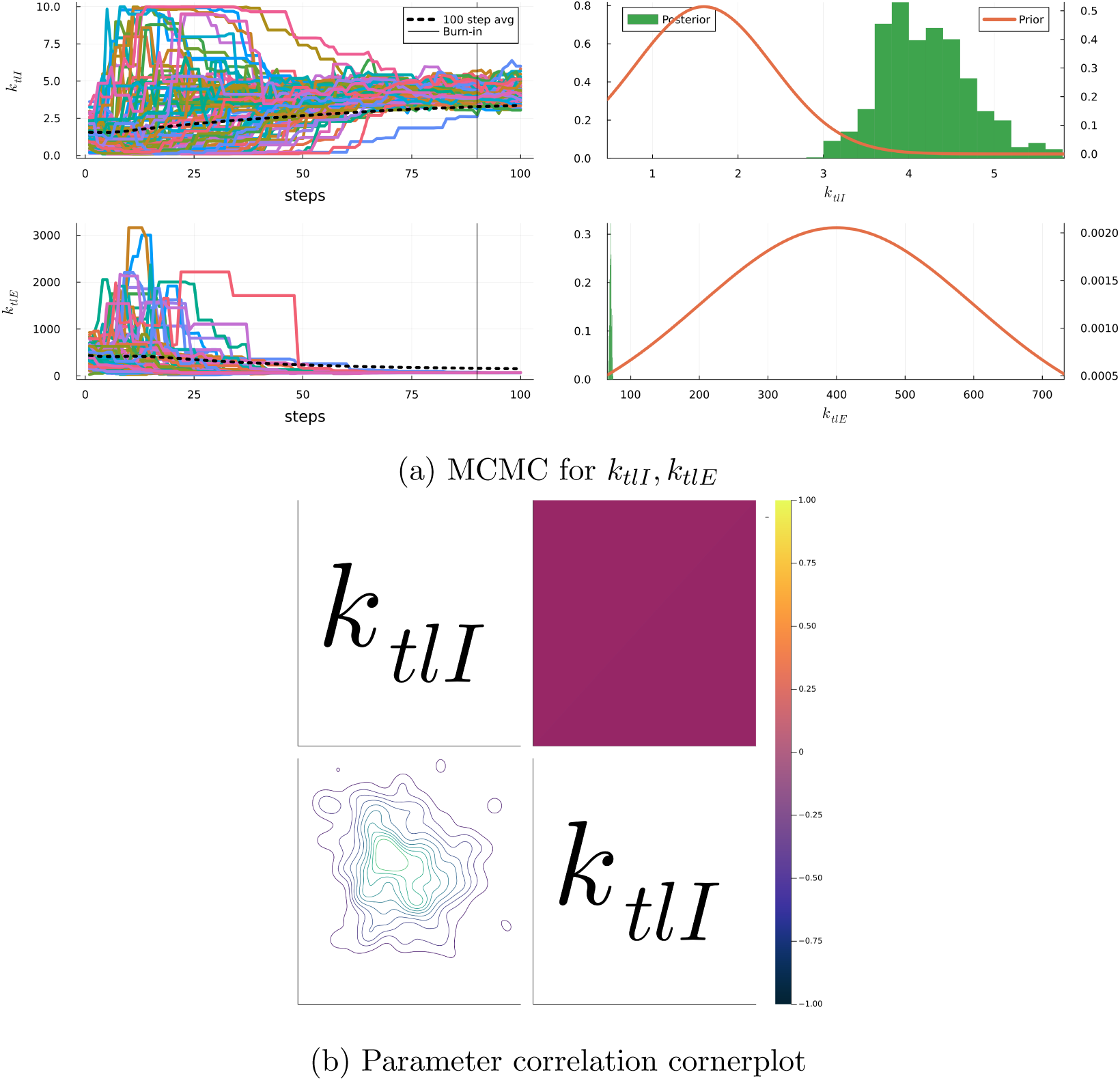
(a) Raw chain trajectories and posterior for *k_tlI_, k_tlE_*, assuming contributions from both. (left) Raw walkers (coloured lines), average over all walkers over the last 100 steps (dashed line), and burn-in steps marked with a vertical line. (right) Density function of the prior distribution (orange, right y-axis), compared to posterior distribution after burn-in (green, left y-axis). (b) Cornerplot of posterior correlations. Contour plot showing posterior correlations (lower left). Correlation in linear-linear (upper left triangle) and log-log space (lower right triangle) (upper right), according to colourbar on the right.

The observed protein translation rate may be obscured by other phenomena influencing the measurable protein content, such as premature translation. This has been observed to be impactful in CFPS, depending on the construct and system.^15,54–56^ We took the possibility of protein truncation into account in a separate model by introducing a truncation probability *p_t_ ∈* [0, 1]. This value quantifies the probability of truncation per amino acid, by implementing the frequency of non-termination

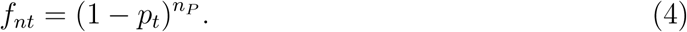

As can be seen, *f_nt_* decreases with *p_t_* and the length of the protein *n_P_* . This is integrated into the model by assuming that the rate of binding of translating complex is split into two populations *c* and *c_t_*, where *c_t_* is translating complex that produces truncated, nonobservable protein, but otherwise behaves identically to *c*, giving

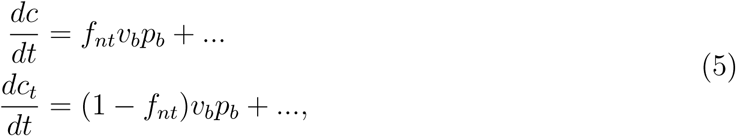

where *p_b_* is the polysome density on the mRNA. With this model expansion, we carried out the same analysis on rate limitations, but inferring the value of *p_t_* alongside *k_tlI_* and *k_tlE_*. This gave a better fit to the data for any of the three combinations (Figs. S11-S13), suggesting that something like protein truncation may indeed be the cause of the observed data, however this would have to be tested experimentally for any conclusive statement.

**Figure S11:**
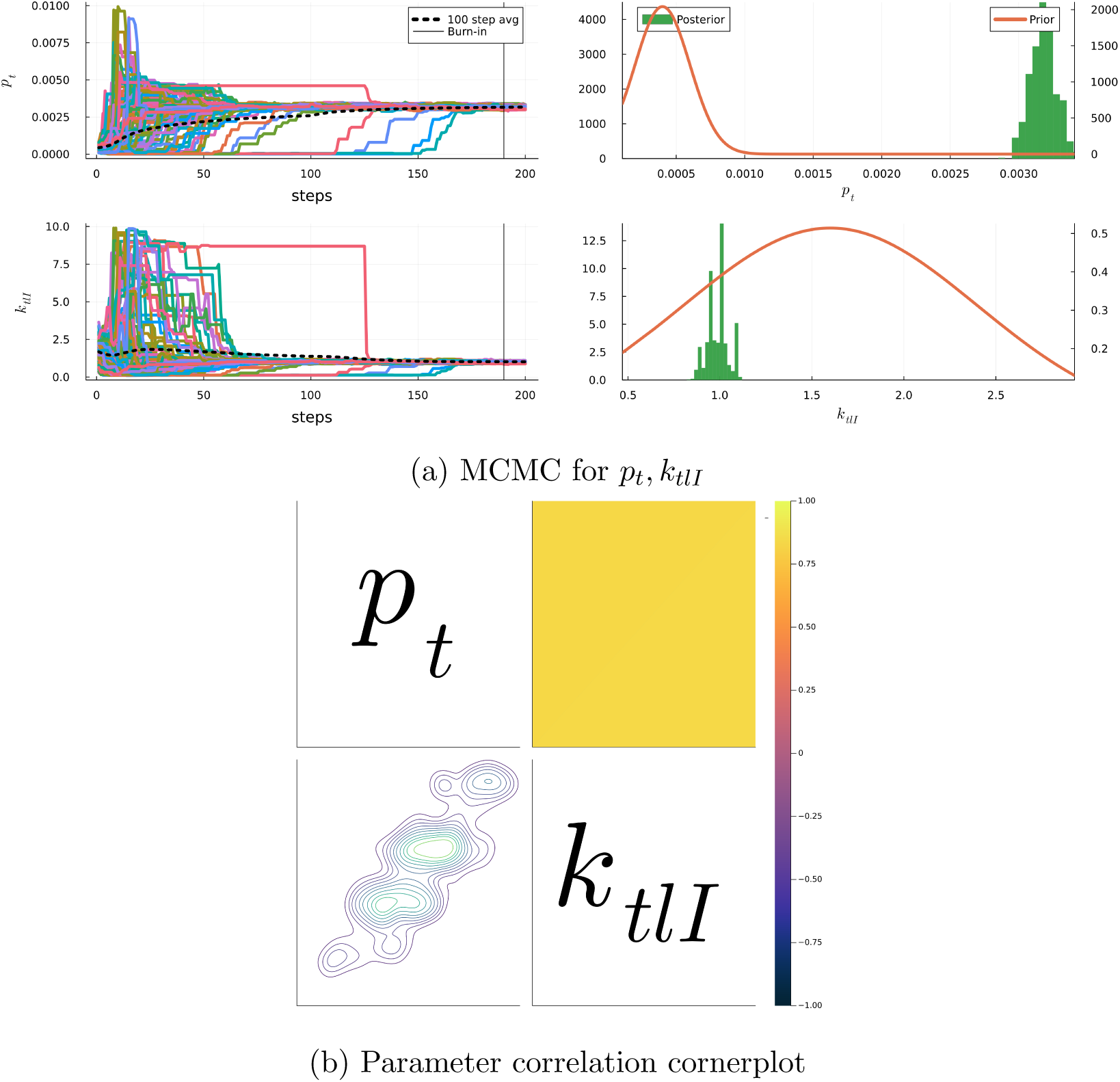
(a) Raw chain trajectories and posterior for *k_tlI_* and *p_t_*, assuming initiation-limited translation with protein truncation. (left) Raw walkers (coloured lines), average over all walkers over the last 100 steps (dashed line), and burn-in steps marked with a vertical line. (right) Density function of the prior distribution (orange, right y-axis), compared to posterior distribution after burn-in (green, left y-axis). (b) Cornerplot of posterior correlations. Contour plot showing posterior correlations (lower left). Correlation in linear-linear (upper left triangle) and log-log space (lower right triangle) (upper right), according to colourbar on the right.

**Figure S12:**
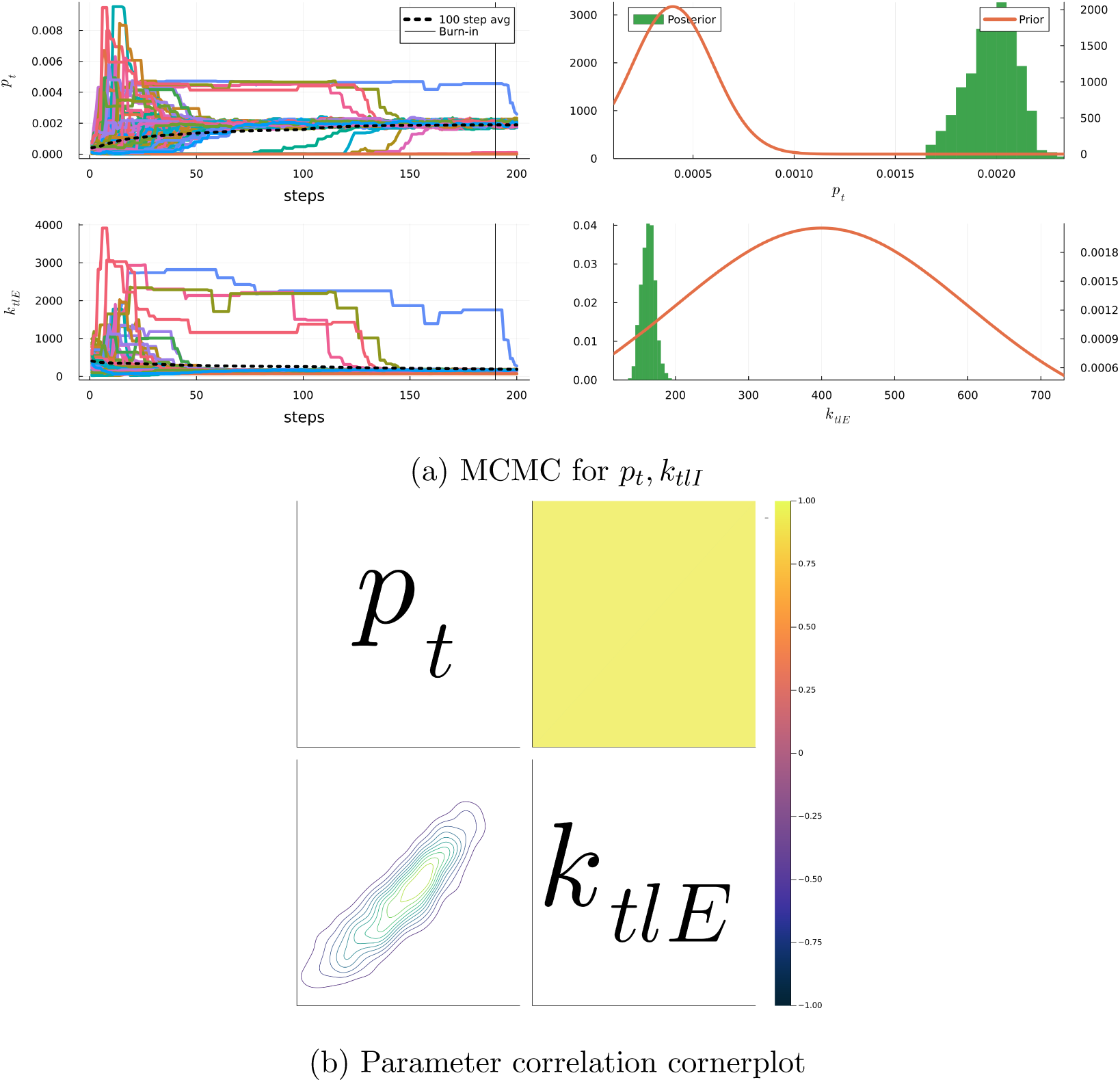
(a) Raw chain trajectories and posterior for *k_tlE_* and *p_t_*, assuming elongation-limited translation with protein truncation. (left) Raw walkers (coloured lines), average over all walkers over the last 100 steps (dashed line), and burn-in steps marked with a vertical line. (right) Density function of the prior distribution (orange, right y-axis), compared to posterior distribution after burn-in (green, left y-axis). (b) Cornerplot of posterior correlations. Contour plot showing posterior correlations (lower left). Correlation in linear-linear (upper left triangle) and log-log space (lower right triangle) (upper right), according to colourbar on the right.

**Figure S13:**
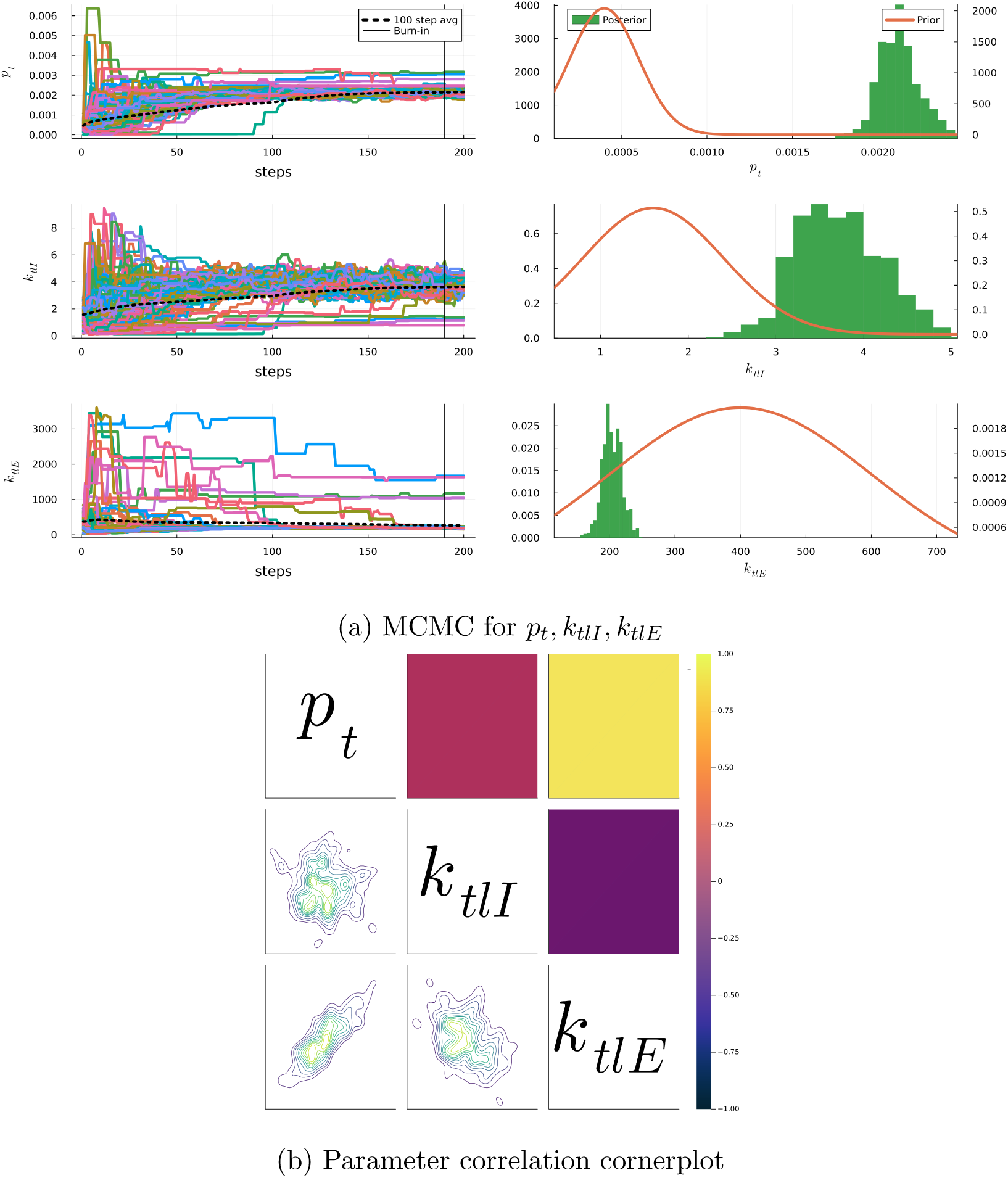
(a) Raw chain trajectories and posterior for *k_tlI_, k_tlE_, p_t_*, assuming translation limited by both and protein truncation. (left) Raw walkers (coloured lines), average over all walkers over the last 100 steps (dashed line), and burn-in steps marked with a vertical line. (right) Density function of the prior distribution (orange, right y-axis), compared to posterior distribution after burn-in (green, left y-axis). (b) Cornerplot of posterior correlations. Contour plot showing posterior correlations (lower left). Correlation in linear-linear (upper left triangle) and log-log space (lower right triangle) (upper right), according to colourbar on the right.

### Batch-to-batch variability inference

We processed the data from all experimental assays as outlined in the methods. We then trimmed each timeseries to the same time period, 0-460 min, to remove the bulk of the data after termination of the reaction. A few reactions had to be trimmed further, due to technical issues during the assay. The affected reactions were all part of the ribosome titration assay. S150_1_ with 0.81 µM ribosomes had to be trimmed to 230 min, with 0.59 µM to 320 min, with 0.33 µM to 420 min, S150_2_ with 0.81 µM to 360 min and S150_3_ with 0.81 µM to 420 min. This removal of timepoints will have slightly decreased the weight of the ribosome assay.

We then scaled each assay via its reference experiment to the yield of the DNA assay reference experiment, by calculating the relative difference between the reference experiment of the other assays to the DNA reference experiment and scaling them up to match (Fig. S14). We did this after reasoning that the reference reactions between experiments should be identical in theory and hence such a normalisation may remove experimental variation we are not interested in modelling.

We fed the normalised data into the Bayesian inference pipeline and scaled the parameters of interest to a mean of 1, only scaling them back up for prediction purposes. Inference ran until the posterior had visually converged, which took 100 walkers around 3000 steps for S150_1_ and 2000 steps for S150_2_. S150_3_ failed to converge all walkers within 20,000 steps.

We therefore restarted inference with the final parameter values from the S150_2_ chain as starting values and the chain finally converged after another 2000 steps (Fig. S15, Tab. S4). We sampled posteriors from the last 100 steps of each chain (Fig. S16).

We calculated parameter correlations as previously explained. On top, we calculated the mean of the absolute correlation coefficients for each batch as 0.22 for S150_1_, 0.24 for S150_2_ and 0.34 for S150_3_ (Fig. S17).

**Table S4:**
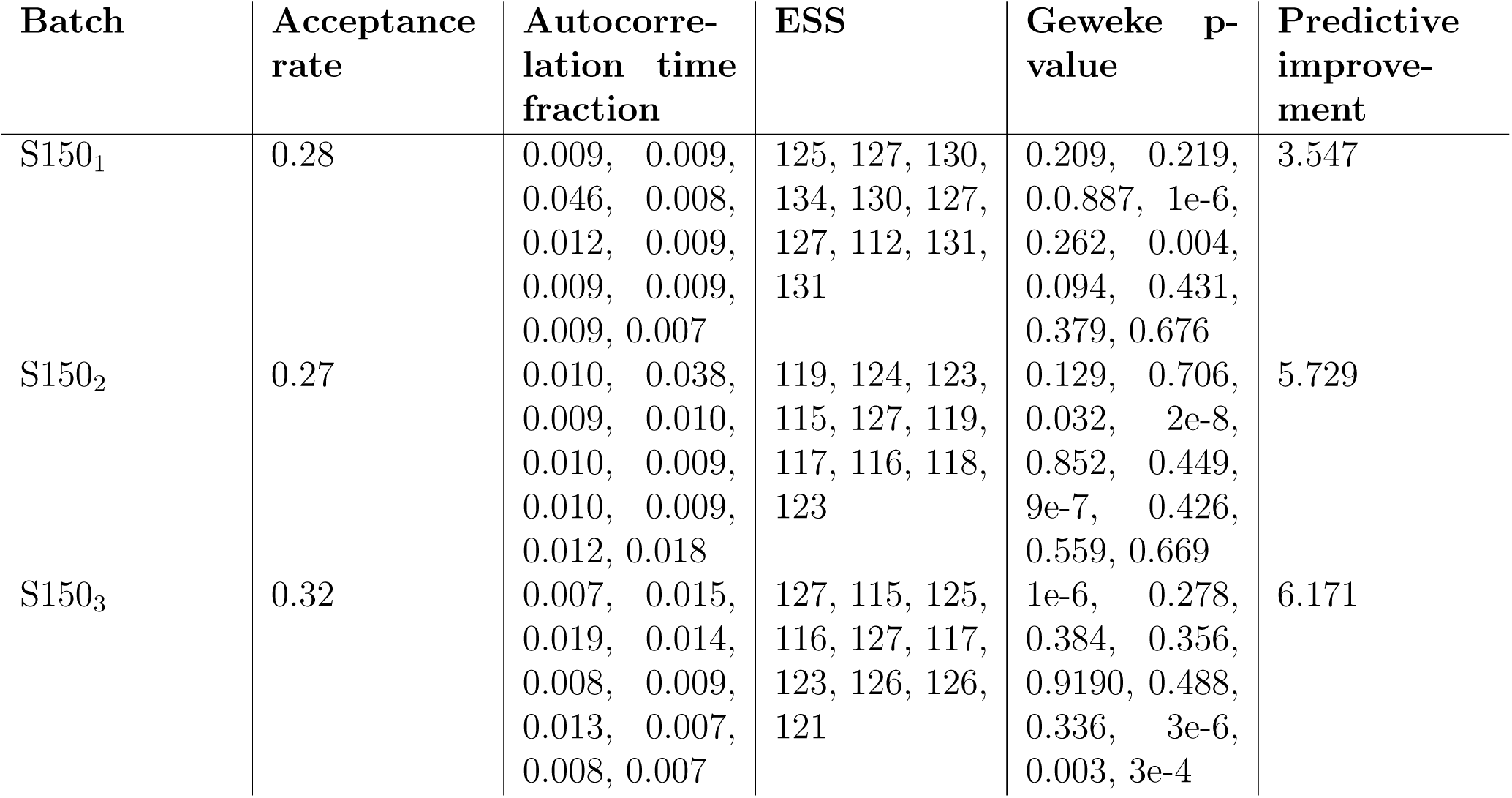
Convergence metrics for batch-to-batch variability study. Inferred parameters were in that order *e*_0_, *k_tx_*, *k_tl_*, *k_d_*, *k_m_*, *k_de_*, *K_tx_*, *K_tl_*, *q_x_*, *q_l_*.

| **Batch** | **Acceptance rate** | **Autocorre-lation time fraction** | **ESS** | **Geweke p-value** | **Predictive improve-ment** |
|---|---|---|---|---|---|
| S150<sub>1</sub> | 0.28 | 0.009, 0.009, 0.046, 0.008, 0.012, 0.009, 0.009, 0.009, 0.009, 0.007 | 125, 127, 130, 134, 130, 127, 127, 112, 131, 131 | 0.209, 0.219, 0.0.887, 1e-6, 0.262, 0.004, 0.094, 0.431, 0.379, 0.676 | 3.547 |
| S150<sub>2</sub> | 0.27 | 0.010, 0.038, 0.009, 0.010, 0.010, 0.009, 0.010, 0.009, 0.012, 0.018 | 119, 124, 123, 115, 127, 119, 117, 116, 118, 123 | 0.129, 0.706, 0.032, 2e-8, 0.852, 0.449, 9e-7, 0.426, 0.559, 0.669 | 5.729 |
| S150<sub>3</sub> | 0.32 | 0.007, 0.015, 0.019, 0.014, 0.008, 0.009, 0.013, 0.007, 0.008, 0.007 | 127, 115, 125, 116, 127, 117, 123, 126, 126, 121 | 1e-6, 0.278, 0.384, 0.356, 0.9190, 0.488, 0.336, 3e-6, 0.003, 3e-4 | 6.171 |

**Figure S14:**
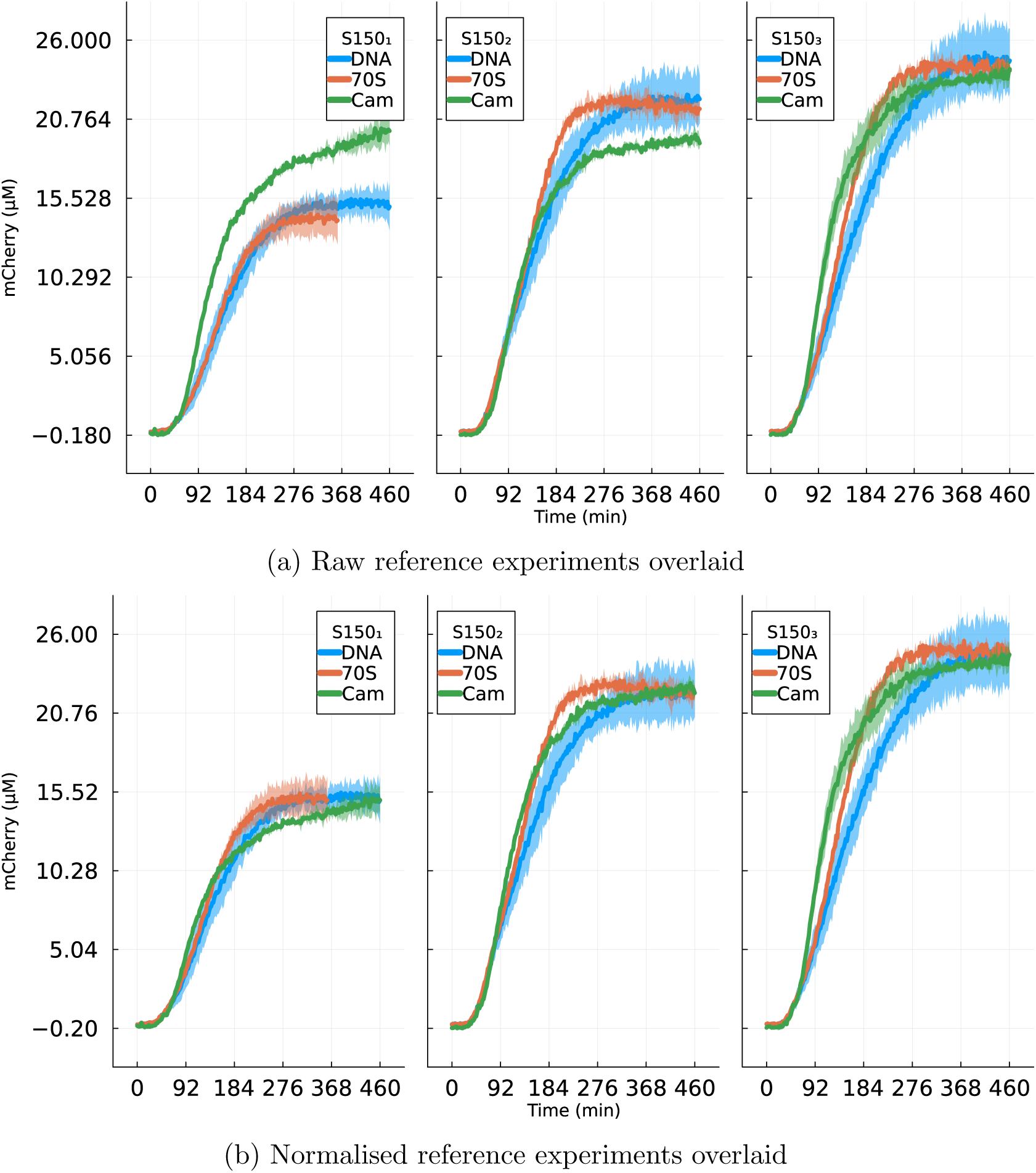
Reference experiment alignment. (a) The raw reference reactions for each assay and batch overlaid. Solid line shows the average and shaded area one standard deviation over triplicates (DNA and 70S titrations) or duplicates (chloramphenicol addition). (b) Each reference reaction normalised to the maximum value of the DNA reference reaction.

**Figure S15:**
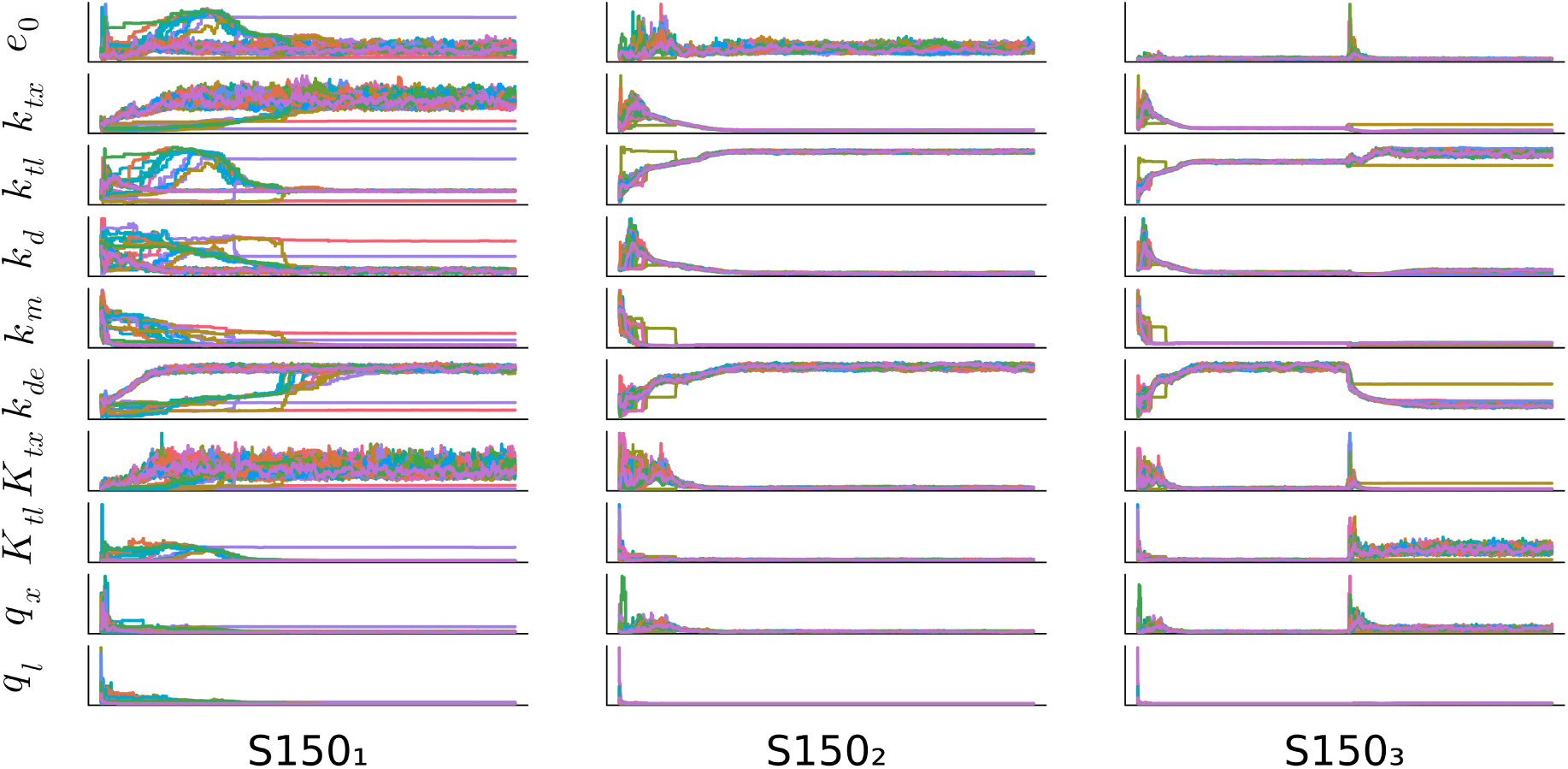
Raw chain trajectories for inference of *e*_0_, *k_tx_*, *k_tl_*, *k_d_*, *k_m_*, *k_de_*, *K_tx_*, *K_tl_*, *q_x_* and *q_l_* using data from extract batches S150_1_-S150_3_. We ran each inference with 100 walkers, starting from the same prior distributions. Inference of parameters for S150_3_ starting from priors had convergence issues, so we restarted inference on the final chain state from S150_2_ instead.

**Figure S16:**
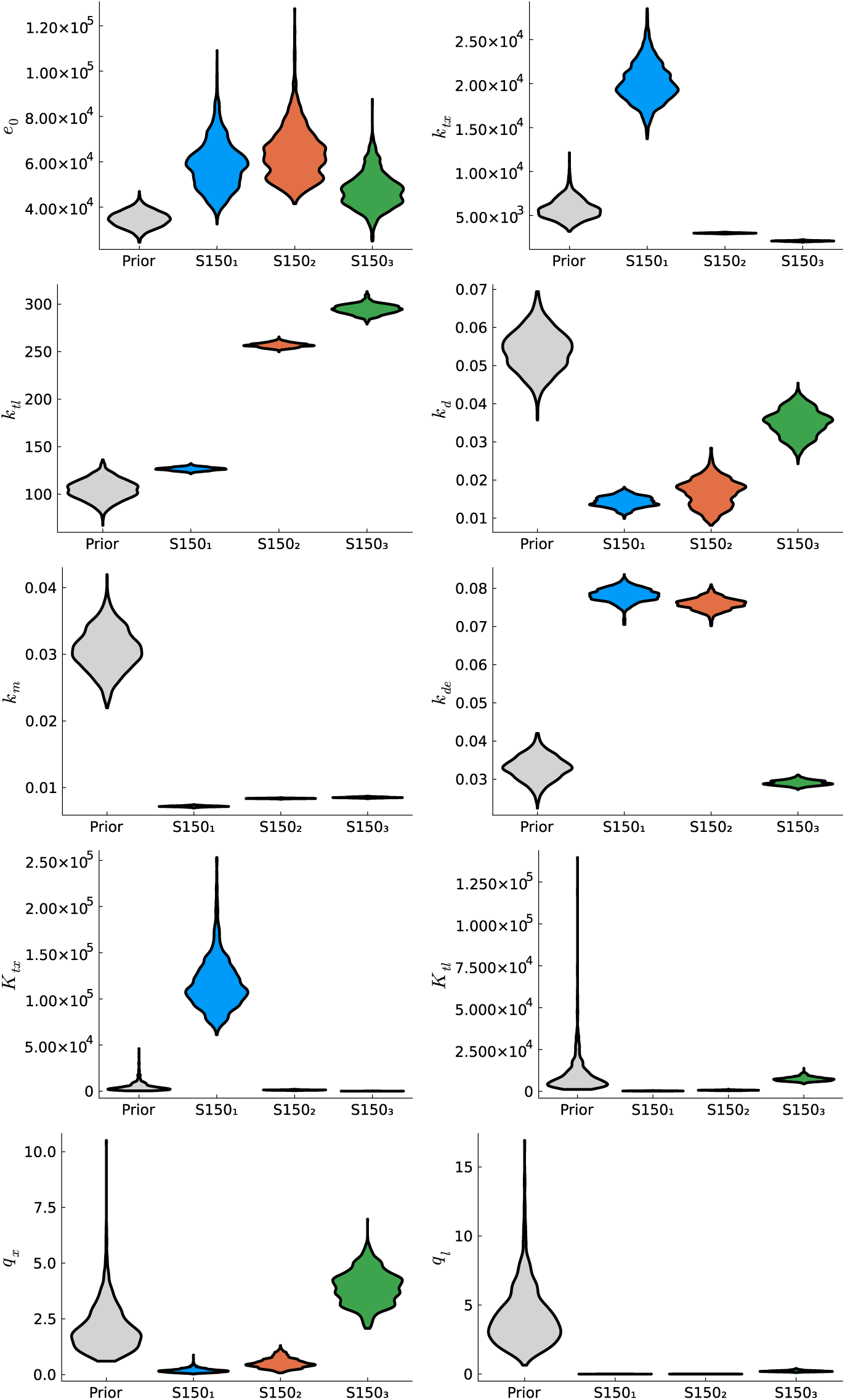
Violin plots of prior and posterior distributions for each batch.

**Figure S17:**
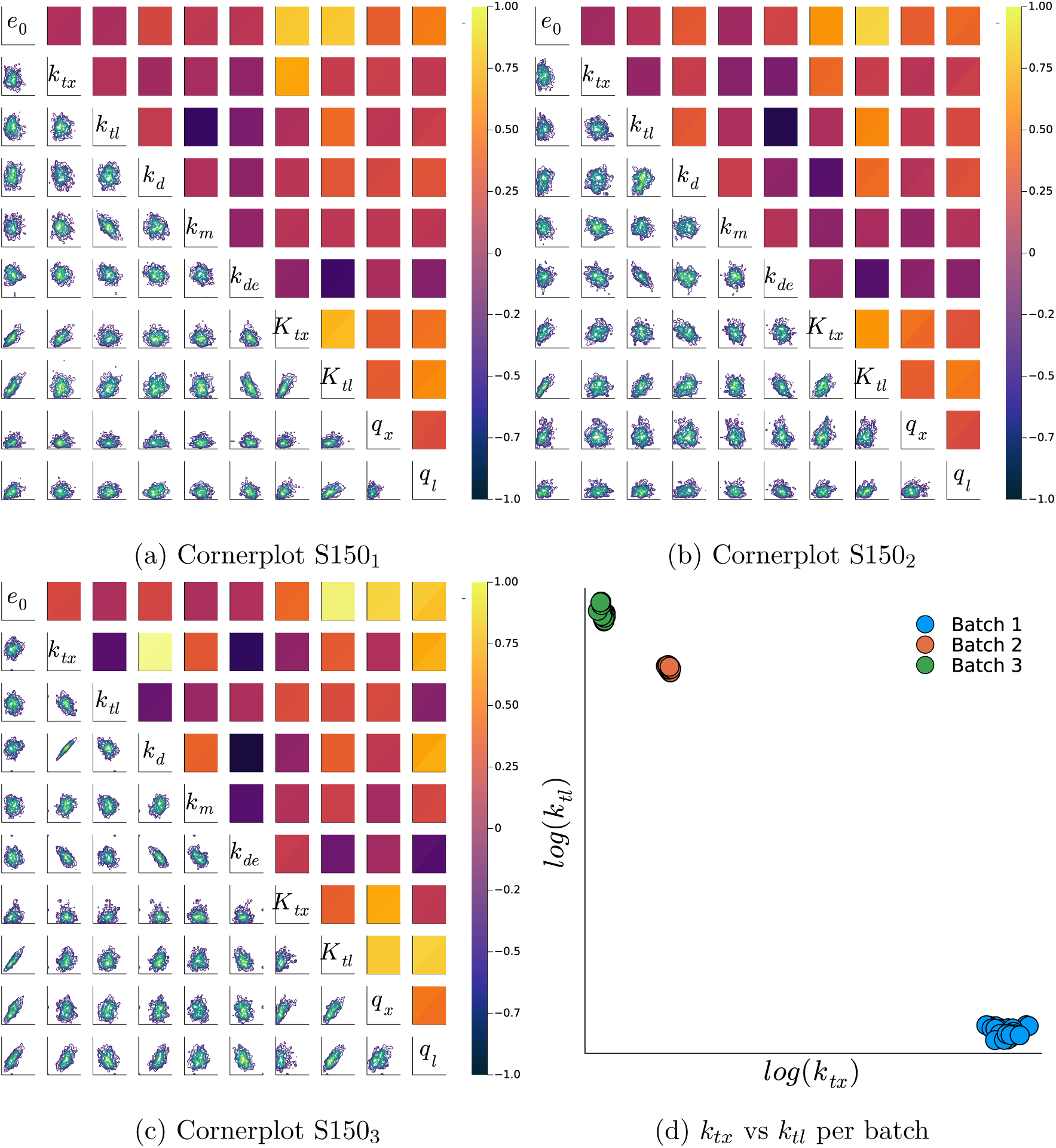
Cornerplots of posterior correlations for inferences on data from S150_1_-S150_3_ (a-c). Contour plot showing posterior correlations (lower left). Correlation in linear-linear (upper left triangle) and log-log space (lower right triangle) (upper right), according to colourbar on the right. (d) 100 samples from posterior parameter distributions of *k_tx_* and *k_tl_* plotted against each other in log-log space for all three batches.

### Calibration curves

To make readings from different platereaders comparable and turn platereader RFU output into tangible micromolar concentration units, we produced calibration curves using purified mRNA and protein. We prepared dilution series of the Broccoli aptamer in extract buffer supplemented with DFHBI-1T at 3-200 ng µL*^−^*^1^ and of mCherry in extract buffer at 4-2000 µg mL*^−^*^1^ and measured them in triplicate on the platereaders at 37 *^◦^*C, after allowing the fluorescence signal to stabilise. We then plotted the resulting data in RFU against the corresponding concentration and fitted a linear function by weighted least squares (Fig. S18). The function coefficients for conversion of Broccoli RFU into mass concentration were

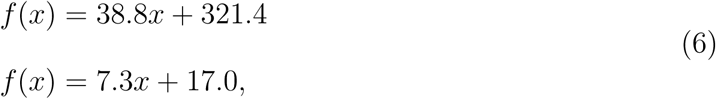

for Fluostar and Biotek, respectively. For mCherry the respective functions were

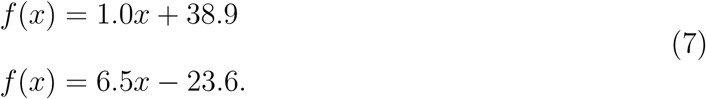

We subsequently converted the relations into molar concentrations by means of the respective molecule molar weights.

**Figure S18:**
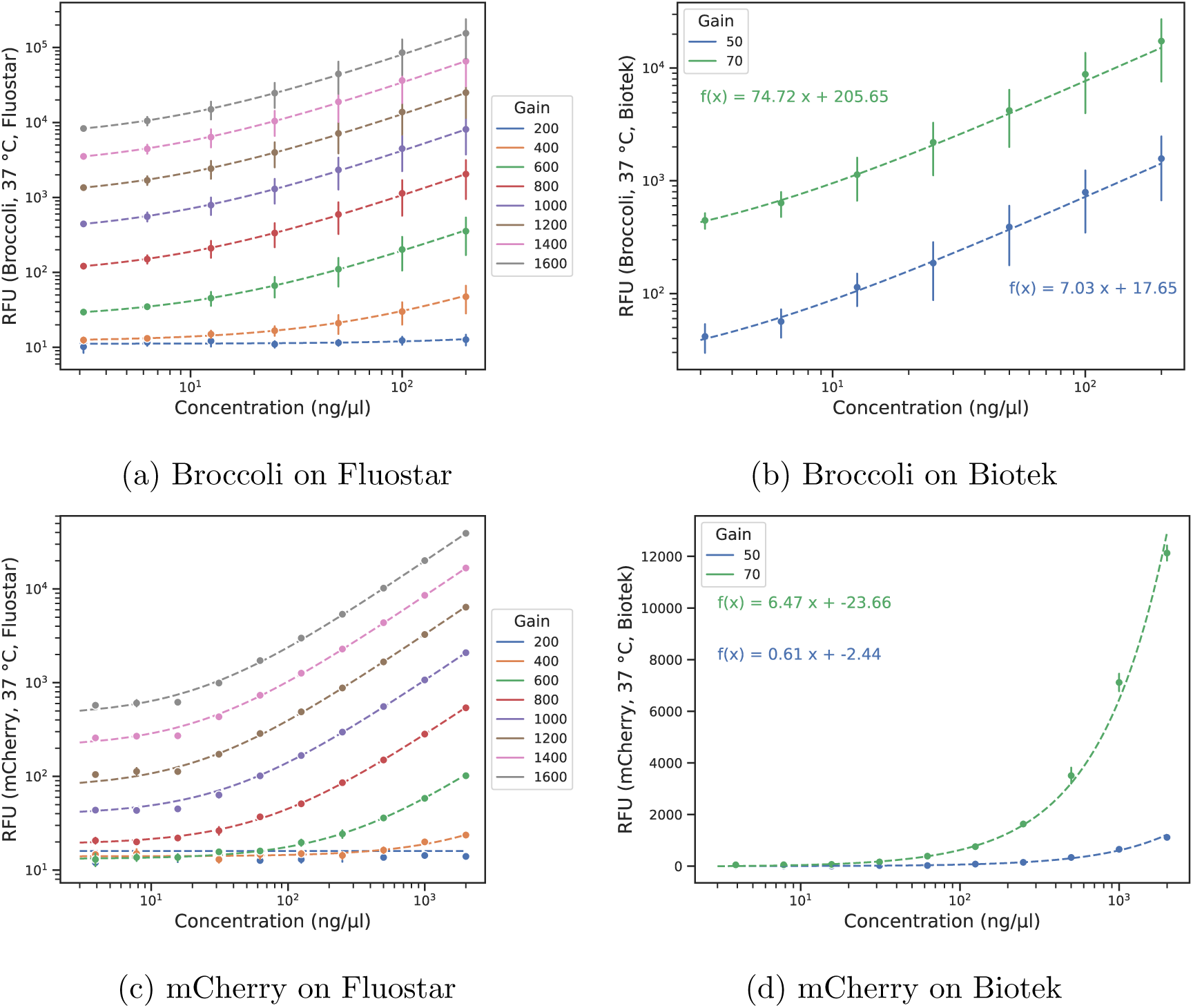
Calibration curves. (a) Calibration of Broccoli on the Fluostar platereader at varying gain settings. We used only gain 1000 later on. (b) Calibration of Broccoli on the Biotek platereader at two gain settings. We used only gain 50 later on. (c) Like (a) but for mCherry. (d) Calibration of mCherry on the Biotek platereader at two gain settings. We used only gain 70 later on. Each concentration was measured in triplicate, while Broccoli measurements were performed from two batches of purified RNA. This gave an estimate for the average signal (solid line) alongside one standard deviation (shaded area).

### Other

**Figure S19:**
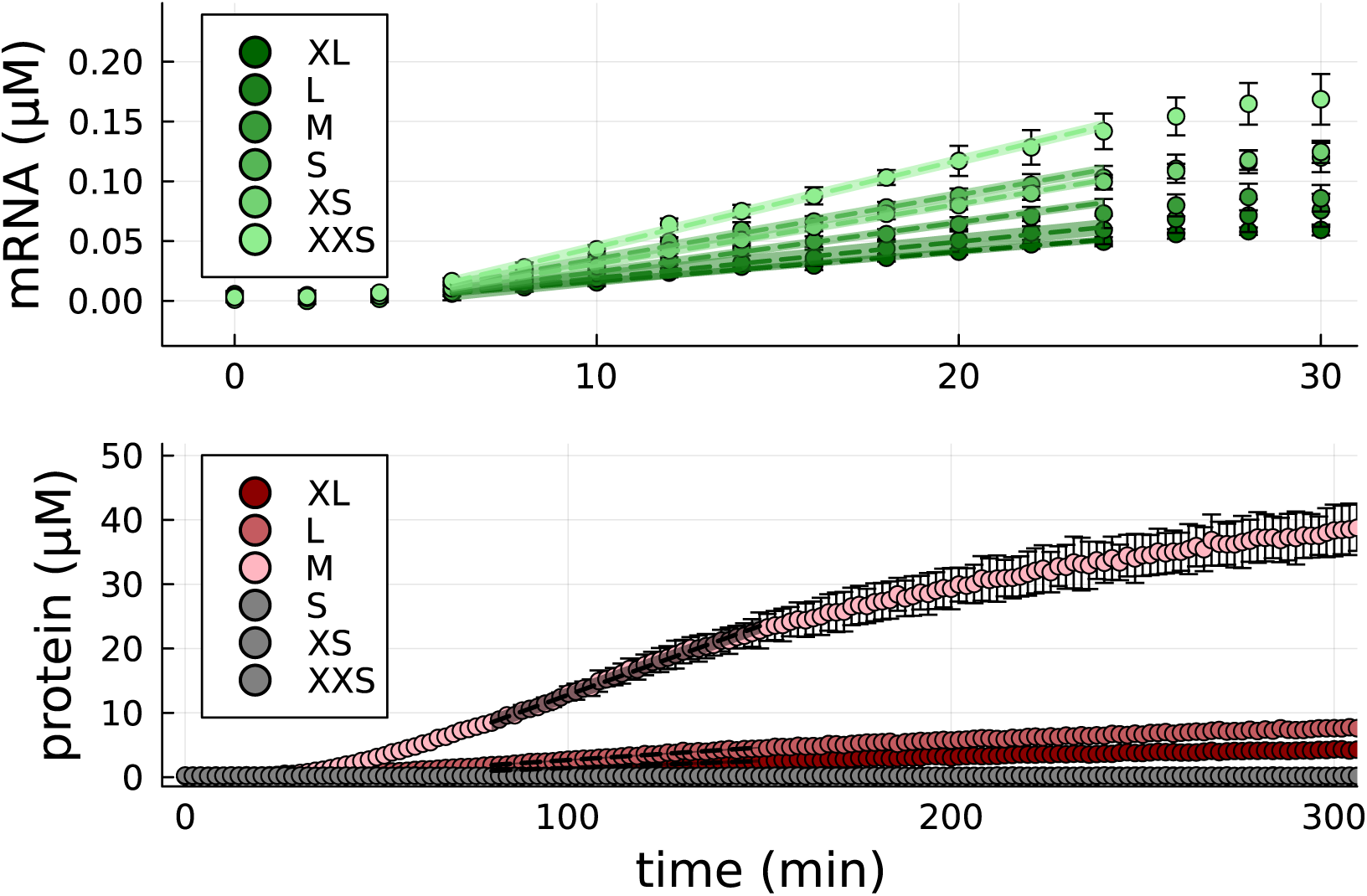
Linear fits for transcription and translation. We analysed both sets of data analogously. We extracted transcription data (top) between 6-25 min, and translation data (bottom) between 80-150 min. We then calculated weighted least squares fits. The shading corresponds to the length of the gene. Grey data in the translation graph corresponds to the three shortest genes, which do not produce a mature mCherry. Circles denote the average and errorbars one standard deviation over triplicates.

**Figure S20:**
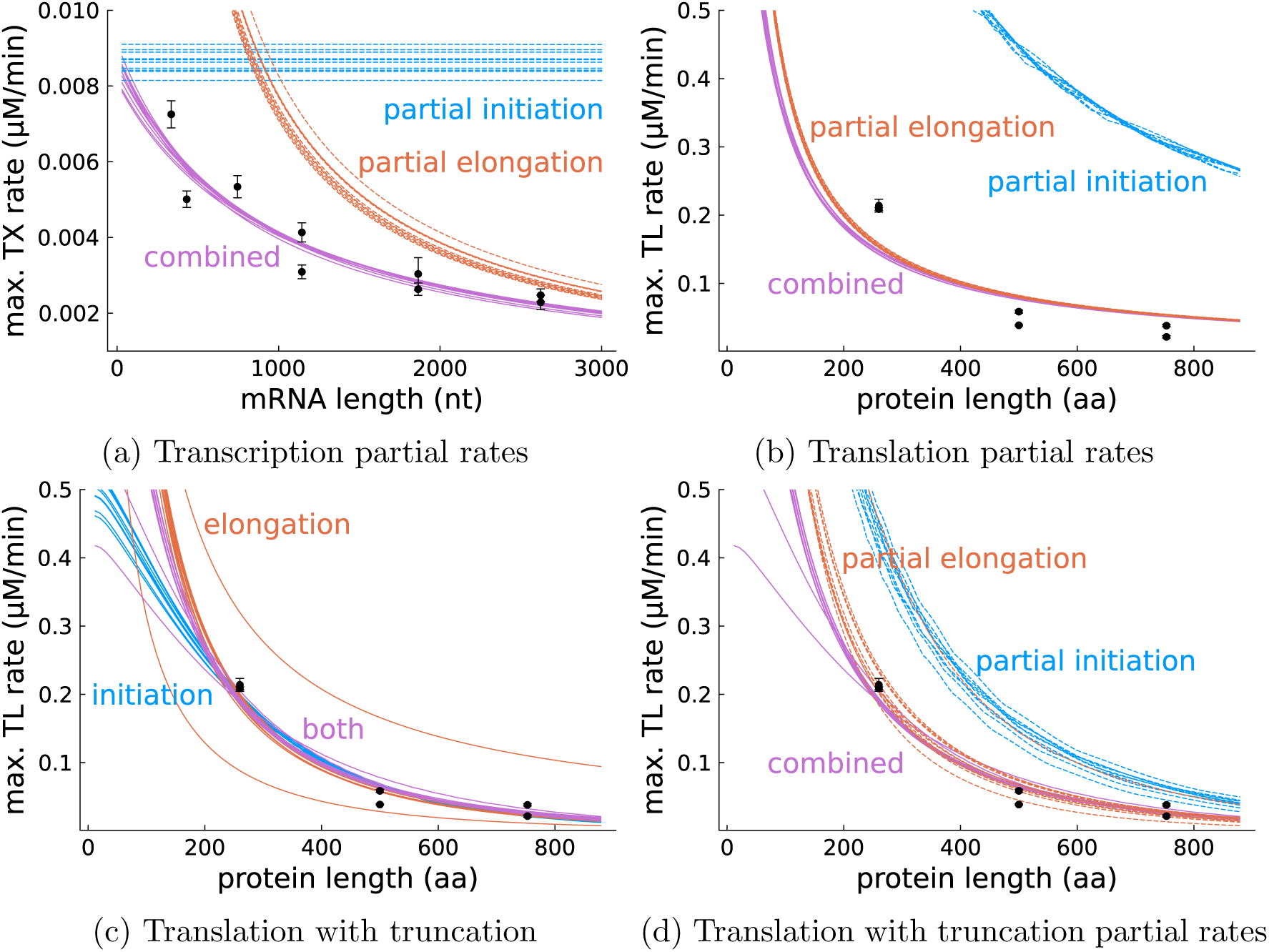
Prediction of transcription and translation rates after model inference under various hypotheses. (a) Transcription inferred under the hypothesis that both initiation and elongation contribute to the combined rate (magenta). The contributions of initiation (blue, dashed) and elongation (orange, dashed) are plotted individually and together make up the combined curve. (b) Same as (a) but for translation. (c) Prediction of translation rates inferred under the assumption of protein truncation. The plot is analogous to Fig. 3E. (d) Same as (a) and (b) but for (c).

**Figure S21:**
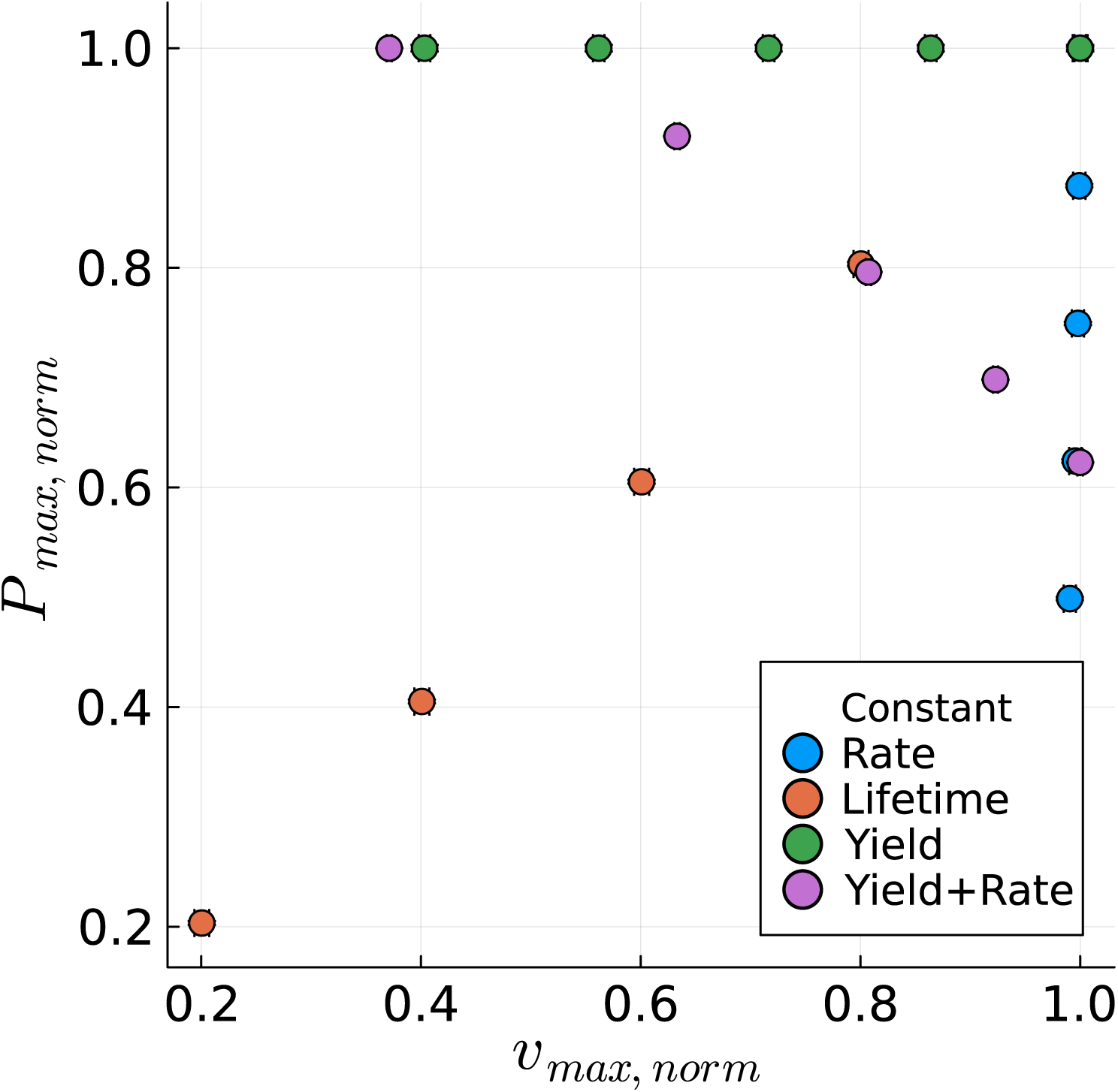
Four trajectories along the Yield-Rate space are conceivable: A system that is rate-limited can achieve varying yield by adjusting the substrate availability. A system that is yield-limited can be varied in its rate. A lifetime-limited system achieves a yield proportional to its rate. A system, where the lifetime decreases with system activity, e.g. if a more active enzyme loses activity more quickly than a less active one, can theoretically display the orthogonal diagonal to the lifetime-limited system.

**Figure S22:**
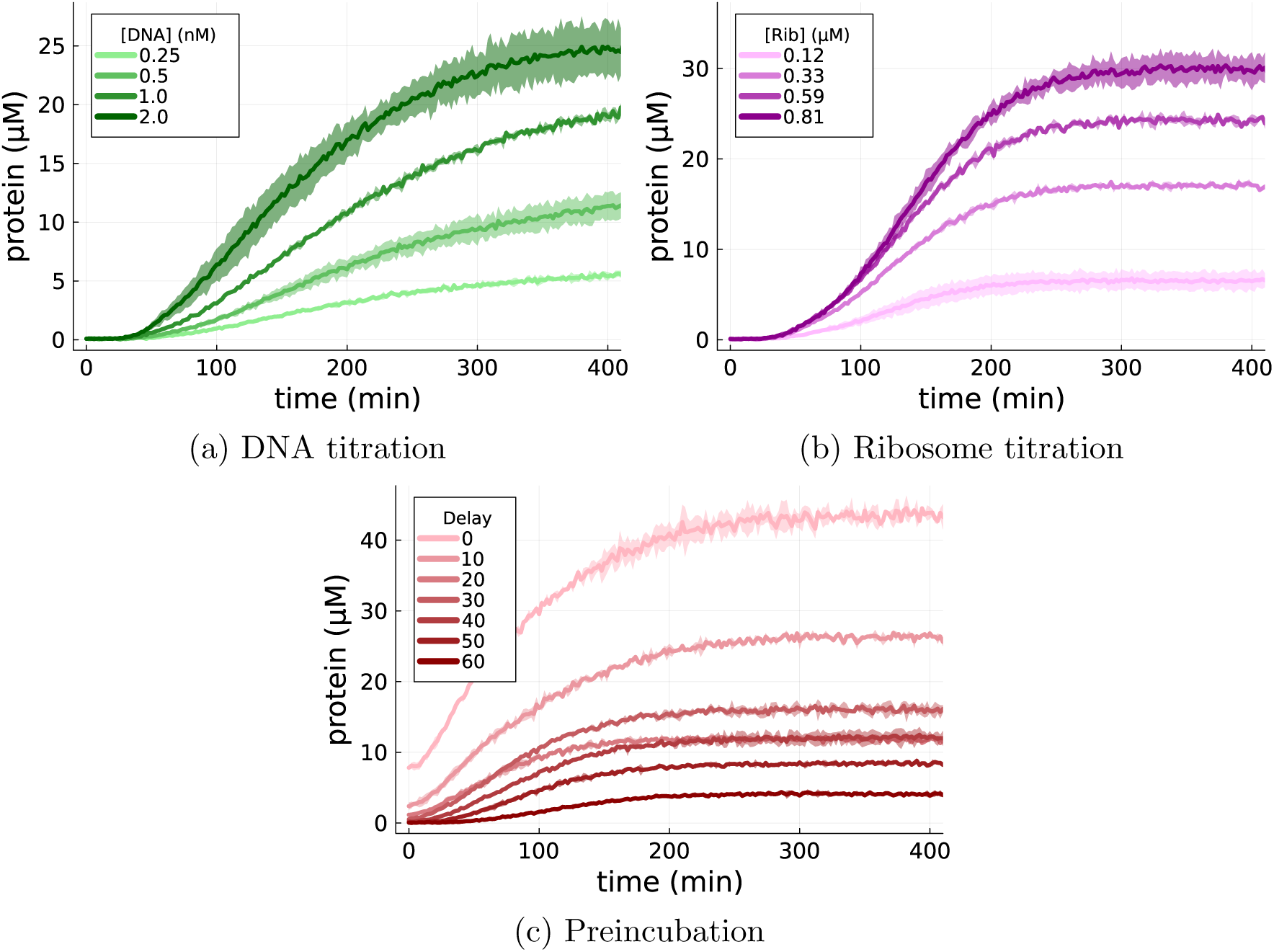
Experimental data used for populating the rate-yield plot to determine the termination mechanism of our system. (a) Reactions with varying concentrations of template DNA. (b) Reactions with varying concentrations of ribosomes. (c) Reactions preincubated for varying time periods at 37 *^◦^*C without template DNA before addition of template DNA. Solid lines show the average and shaded areas one standard deviation over triplicates (a, b) or duplicates (c).

**Figure S23:**
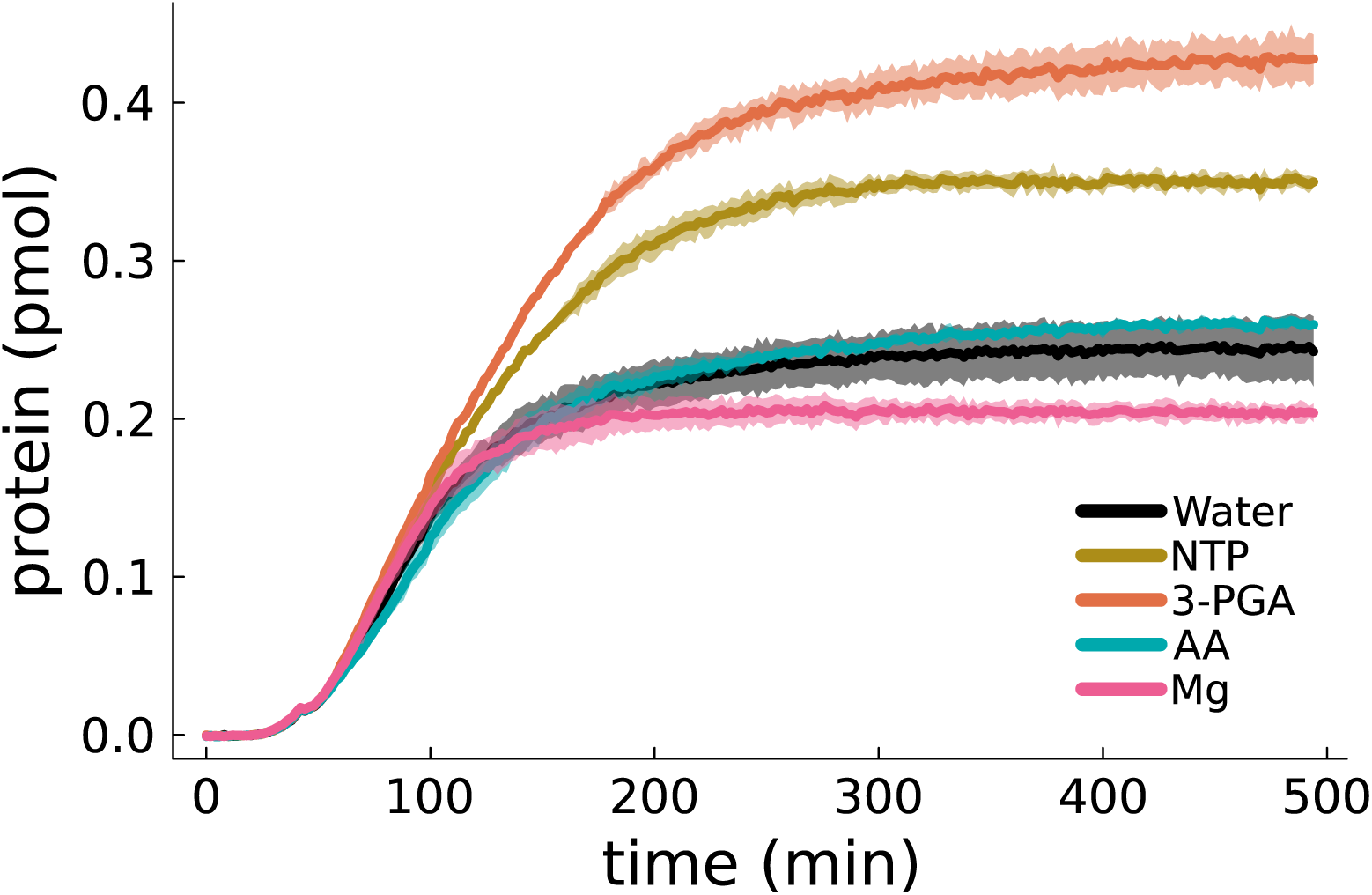
Reactions spiked with a selection of energy solution components, to identify the first limiting factor. We prepared identical default reactions simultaneously. After 40 min, we added fresh NTPs, 3-PGA, Mg-Glutamante or amino acids to separate reactions in duplicate. We added H_2_O as a negative control. Solid lines denote the average and shaded area one standard deviation over duplicates.

**Figure S24:**
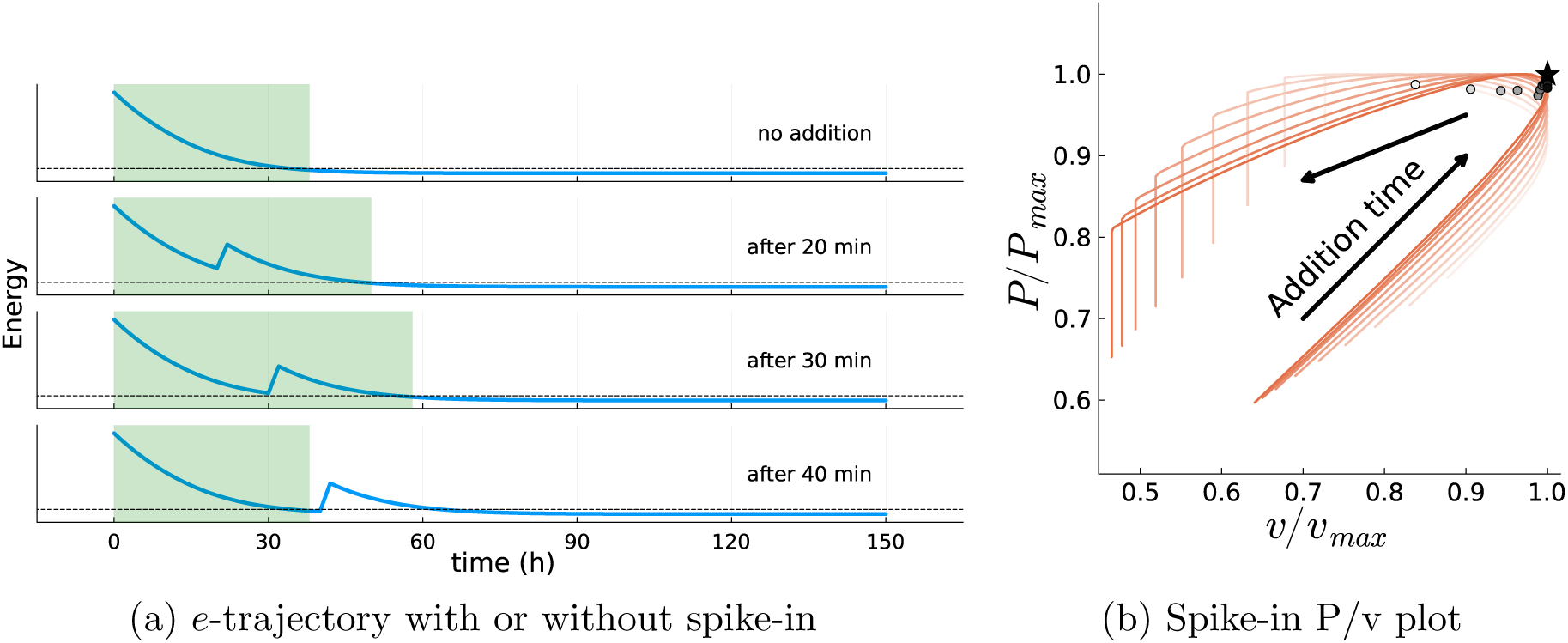
Energy trajectory approaching the depletion limit. (a) Plotted are four simulations of the model assuming *k_de_* = 0.06 min*^−^*^1^ with spike-in of energy never (top) or after 20, 30 or 40 minutes. The dotted line shows the arbitrary depletion limit at 2 mM. The green box indicates the time the system continuously spent at full activity due to energy abundance. Addition of energy after 20 and 30 minutes prolongs the interval of full activity, while addition after 40 minutes is past the depletion point. (b) Model simulations for spike-in benefit at varying time points and degradation rate constants with the time point of energy depletion marked with a circle for each curve.

**Figure S25:**
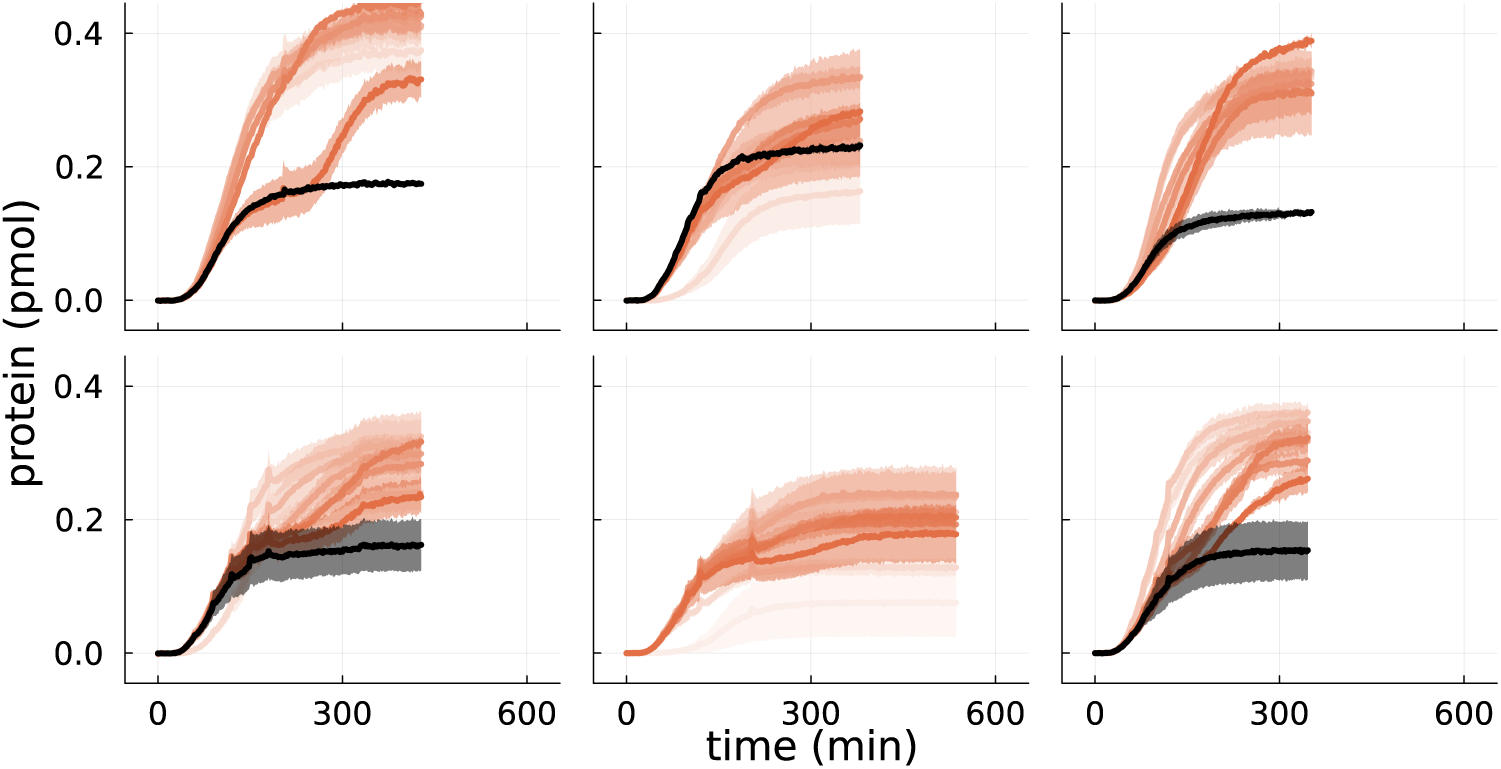
Raw reaction traces for all performed 3-PGA spike-in experiments under three different conditions. (left column) 30 mM 3-PGA at 2 nM DNA every 10 min (top) and every 30 min (bottom). (middle column) 30 mM 3-PGA at 5 nM DNA every 20 min (both). (right column) 15 mM 3-PGA at 2 nM DNA every 10 min (top) and every 20 min (bottom). Solid lines show the average and shaded areas one standard deviation over duplicates.

**Figure S26:**
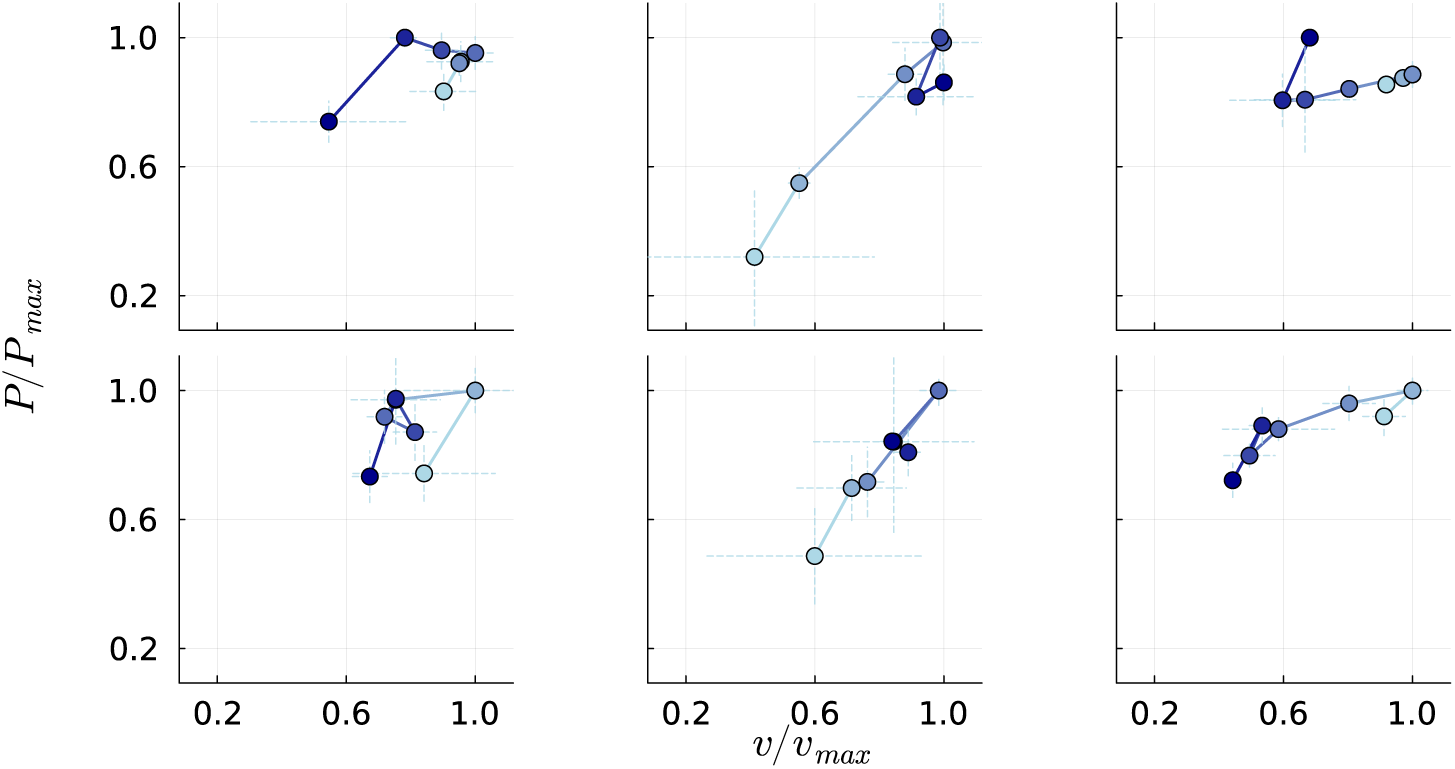
Rate-yield plots for all performed 3-PGA spike-in experiments under three different conditions. (left column) 30 mM 3-PGA at 2 nM DNA every 10 min (top) and every 30 min (bottom). (middle column) 30 mM 3-PGA at 5 nM DNA every 20 min (both). (right column) 15 mM 3-PGA at 2 nM DNA every 10 min (top) and every 20 min (bottom).

**Figure S27:**
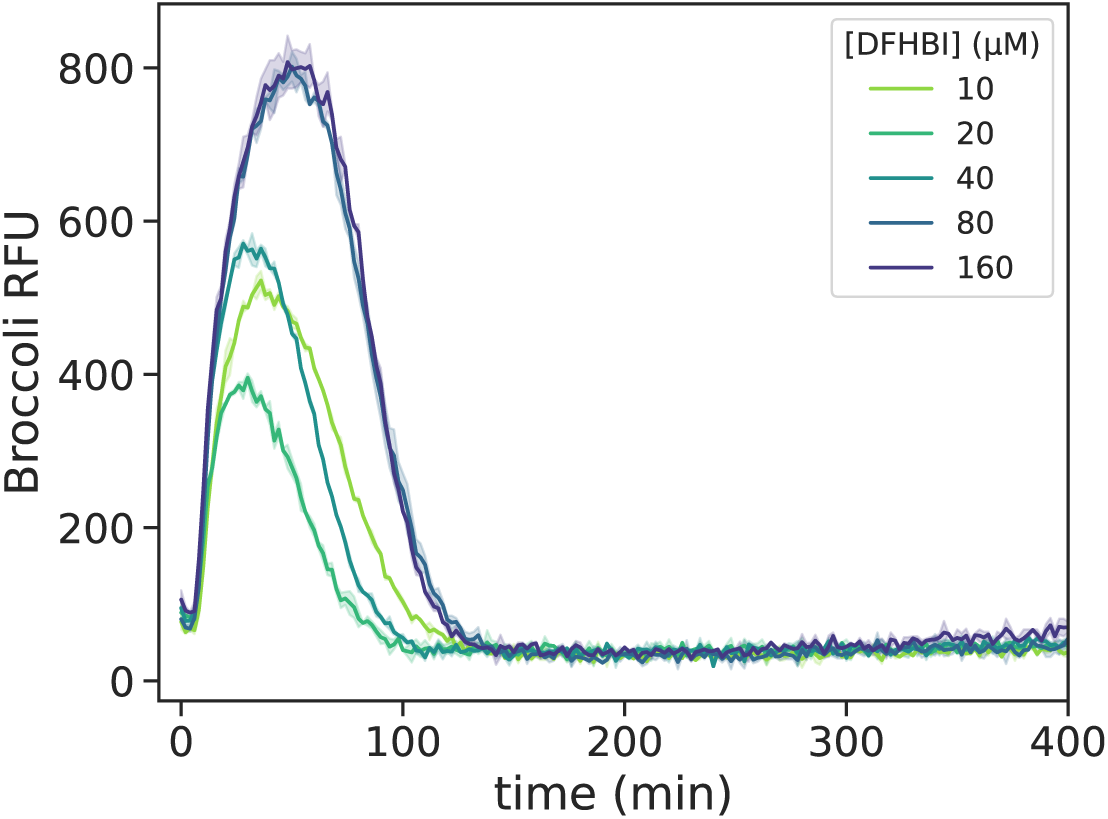
**DFHBI-1T titration** in an S30 reaction with supplemented maltodextrin shows longer and higher peaks when more dye is added. However the initial timepoints of all trajectories align, indicating that the dye is not limiting then and that therefore the readout is reliable.

